# Cooperative electrolyte-PEG interactions drive the signal amplification in a solid-state nanopore

**DOI:** 10.1101/2021.11.01.466478

**Authors:** Chalmers C. Chau, Fabio Marcuccio, Dimitrios Soulias, Martin A. Edwards, Sheena E. Radford, Eric W. Hewitt, Paolo Actis

## Abstract

Nanopore systems have emerged as a leading platform for the analysis of biomolecular complexes with single molecule resolution. However, the analysis of several analytes like short nucleic acids or proteins with nanopores represents a sensitivity challenge, because their translocation lead to small signals difficult to distinguish from the noise. Here, we report a simple method to enhance the signal to noise ratio in nanopore experiments by a simple modification of the solution used in nanopore sensing. The addition of poly-ethylene glycol (PEG) and the careful selection of the supporting electrolyte leads to large signal enhancement. We observed that the translocation dynamics are in good agreement with an established method that uses the lattice energy of an electrolyte to approximate the affinity of an ion to PEG. We identified CsBr as the optimal supporting electrolyte to complement PEG to enable the analysis of dsDNA at 500 kHz bandwidth, and the detection of dsDNA as short as 75 bp.

## INTRODUCTION

Nanopore technology is enabling the analysis of biological macromolecules with single-molecule resolution (1, 2). In nanopore experiments, individual molecules are driven through a nanopore under the influence of an electric field, causing a temporary modulation in the conductance within the pore produced by a combination of the geometrical exclusion of solution, ion concentration polarization and additional charges brought by the analyte itself (3, 4). The magnitude and duration of this momentary change in the ionic current reflects the properties (e.g., size, shape, charge) and the translocation dynamics of the molecule (5-20). There are two classes of nanopores: biological nanopores and solid-state nanopores. The former is based on protein nanopores that have been employed to great success for the real-time sequencing of nucleic acids (21, 22). The latter offers an alternative based on inorganic materials that could provide higher signal-to-noise levels, diameter tunability and improved stability (1, 23, 24).

The analysis of short nucleic acids and proteins with nanopores represents a sensitivity challenge, as the translocation of small molecules leads a signal that is difficult to reliably distinguish from the background current (1, 24). While several approaches have been used to address this challenge, they only partially address the whole problem. For example, the best solution to address both of these challenges relies on t using of nanopores of few nm in diameter embedded in nm-thick membranes integrated with custom designed electronics with MHz bandwidth (11, 23). This approach allows for high signal-to-noise ratio (SNR) and sub μs temporal resolution, but it requires access to highly specialized and costly equipment, which are often accessible to only a handful of laboratories worldwide. Alternative approaches rely on the modification of the physical-chemical properties of the solutions used in nanopore experiments. For example, the viscogen glycerol has been added to the bath electrolyte to reduce the speed of the molecular translocations, but at the expense of a reduced SNR (25). Another approach relies on using LiCl as the electrolyte to slow down the translocation of molecules through nanopores, this approach is particularly effective for nucleic acids but it does not increase the SNR (26). Salt gradients can also be used to improve the translocation frequency across a nanopore, but it does not affect the speed nor the current magnitude of the single molecule events (27). Alternatively, the nanopore surface can be chemically modified to slow down the translocation of analytes (28-32) but this method is difficult to generalize as it needs to be tailored for each analyte.

We have recently reported that dissolving the commonly used macromolecular crowding agent - poly(ethylene) glycol (PEG) at a concentration of 50% (w/v) in the bath solution resulted in a pronounced increase in the SNR for the single molecule detection of DNA, globular and filamentous proteins (33). Here, we build upon our discovery to further characterize the role of the molecular weight of PEG and its interactions with the supporting electrolyte. We determine how the electrolyte employed modifies the translocation dynamics of a model analyte (4.8 kb double-stranded DNA (dsDNA)), as characterized by the translocation event current peak magnitude and the dwell time. Our results indicate that a cooperative effect between the electrolyte and PEG is responsible for driving the signal enhancement in a solid-state nanopore. We observed that the translocation dynamics are in good agreement with an established method that uses the lattice energy to approximate the affinity of an ion to PEG. We identified an atypical salt - CsBr as the optimal supporting electrolyte to complement PEG to enable the analysis of dsDNA at 500 kHz bandwidth, and the detection of dsDNA as short as 75 bp, all with a conventional patch clamp amplifier and a model 25 nm diameter solid-state nanopore that can be easily fabricated at the bench within seconds starting from inexpensive glass capillaries.

## RESULTS AND DISCUSSION

In this study we used nanopipettes as a model solid-sate nanopore as they have been used to great success to analyse a wide variety of analytes at the single-molecule level (1, 3, 14, 28). Nanopipettes with a diameter of *ca*. 25 nm (Supporting Figure 1) were filled with a solution of 0.3 nM dsDNA (4.8 kbp; Supporting Figure 2) diluted in 0.1 M KCl. The nanopipette was immersed into a 0.1M bath solution containing 50% (w/v) PEG (molecular weight (MW) varied) (see Supporting Information for details on the preparation of the bath solutions). The concentration of 50% (w/v) was chosen and kept constant across all experiments, as we previously demonstrated that it provided the highest SNR (33). Two Ag/AgCl electrodes, one inside the nanopipette, the other immersed into the bath solution are used to apply the voltage and measure the current. The translocation of a single dsDNA molecule from inside the nanopipette to the bath solution leads to a current enhancing peak (*i.e*. the dsDNA translocation temporarily increases the measured current), and each peak is a single molecule translocation event (3, 4) (Figure 1A). Each translocation event contains two main characteristics: the current peak maxima (the amplitude of the event) and the dwell time (the width of the event).

**Figure 1.**
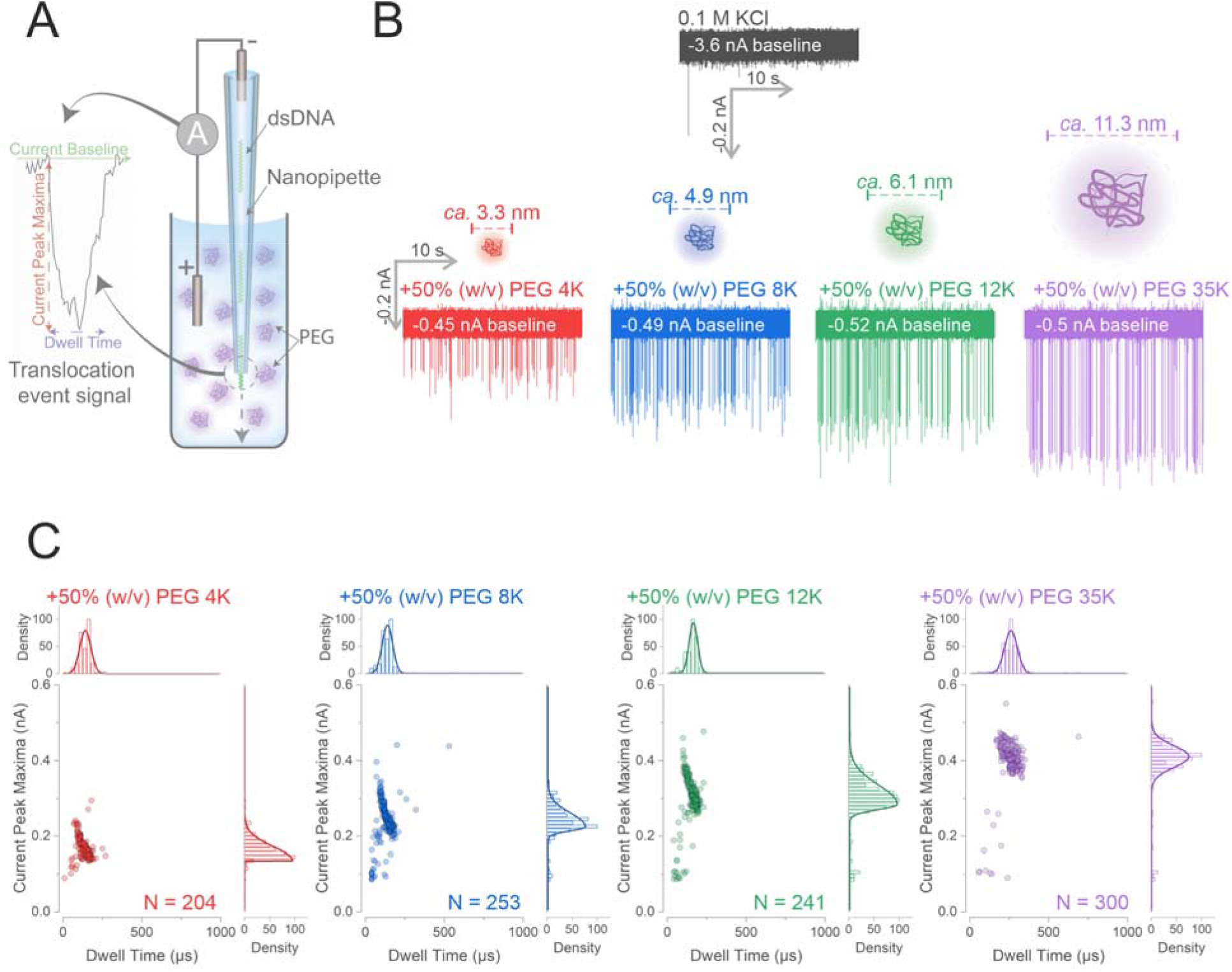
Increasing the MW of PEG enhances the SNR of dsDNA. (A) The experimental set-up schematic, the dsDNA is diluted in 0.1 M KCl to 0.3 nM and used to fill the nanopipette, the nanopipette is then immersed into a 50% (w/v) PEG bath containing 0.1 M electrolyte, a negative voltage is applied from the nanopipette to trigger the translocation of the dsDNA. Each translocation of the dsDNA generates a downward translocation event signal with two main characteristics: the current peak maxima (the change of current level from the current baseline) and the dwell time (the time it requires for the current to return to baseline). (B) The nanopipette was filled with 0.3 nM of 4.8 kbp dsDNA diluted in 0.1 M KCl, the nanopipette was immersed into a 0.1 M KCl bath and an −500 mV was applied to cause the dsDNA to translocate from the nanopipette to the electrolyte bath. The same procedure was repeated in different 0.1 M KCl bath with different PEG molecular weights as stated. The calculated hydrodynamic diameters for the different PEG molecular weight are shown (47). (C) The recorded translocation events in the KCl PEG bath of (B) were used to generate population scatters. All histograms were fitted with a gaussian distributions (for normally distributed data) or bigaussian distribution (for abnormally distributed data). N is the number of events plotted.

Upon the application of −500 mV to the electrode inside the nanopipette, we observed less than 10 translocation events in the 0.1 M KCl electrolyte bath while the addition of PEG increased the number of 4.8 kbp dsDNA molecules detected to 80 (Figure 1B, Supporting Figure 3). We noticed that increasing the MW of PEG lead increase in the current peak magnitude of the single molecule translocation events. The effect of the PEG size can be observed clearly with the population scatter of the translocation events (Figure 1C and Supporting Figure 3) where the population of events shifted to higher peak magnitudes and longer dwell times as we increased the PEG molecular weight (Figure 1C, Supporting Figure 3 and 4A-B). The mean current peak maxima of the dsDNA increased from 0.17±0.02 nA in PEG 4K to 0.40±0.01 nA in PEG 35K, while the dwell time increased from 130±10 μs in PEG 4K to 300±30 μs in PEG 35K, a nearly two-fold increase (Supporting Figure 4A-B). We performed control experiments to ensure that the translocation signals detected were due to the migration of dsDNA from nanopipette to the bath and not to the translocation of PEG itself as this has been reported for biological nanopores (34-41). We did not observe any translocation events when the nanopipette was filled only with 0.1 M KCl and were immersed into the KCl PEG 35K bath (Supporting Figure 5). Since the PEG 35K generated the most significant enhancement in the SNR, it will be used throughout the remaining study unless otherwise stated.

We also measured the shear viscosity and the conductivity of the different molecular weight PEG solutions dissolved in 0.1 M KCl (Supporting Figure 4C-D). A solution of PEG 35K in 0.1 M KCl had a viscosity of nearly 10 Pa·s, which is approximately 100× higher than PEG 4K and 10,000× more viscous than a standard 0.1 M KCl solution. In contrast, the conductivities of all 0.1 M KCl PEG solutions were approximately 1.5 mS/cm, regardless of the MW of PEG. For reference, the conductivity of a 0.1 M KCl solution is approximately 12 mS/cm. The shear viscosity and the conductivity values measured here were in agreement with those reported in the literatures (42, 43). It is interesting to observe that the current peak maxima did not increase by 10× between the PEG 12K and 35K solution despite a nearly a 10× difference in solution viscosity, thus viscosity alone by itself could not fully explain the observed enhancement. Furthermore, these results agreed with our previous observations that the viscogen glycerol did not lead to any current signal enhancement (33). However, it is plausible that the increase in the average dwell time observed in PEG 35K could be related to the increase in the solution viscosity, as Fologea *et al*. in 2005 observed this effect in a solid-state nanopore system (25). Since the current peak magnitude of translocation events changed drastically when we increased the molecular weight of the crowding agent - PEG, the excluded volume effect (*i.e*. the macromolecular crowding effect) also increased simultaneously as different studies have stated (44-46), indicating that macromolecular crowding may also play a role beside the effects of solution viscosity.

On the other hand, when we analysed the relationship between the molecule count and the applied voltage (Supporting Figure 6), we observed an exponential relationship in a 0.1 M KCl solution, indicating a barrier-limited capturing of the dsDNA. However, in the presence of PEG, the dsDNA capture can be described by the diffusion-limited regime, as reflected by the linear dependence of the molecule count to the applied voltage (Supporting Figure 6) (9, 13, 16). The main difference that leads to either the barrier-limited regime or the diffusion-limited regime is the local ion environment near the nanopore (9, 13, 16). Here, the dsDNA was translocated from a pure KCl environment through the nanopore to a highly crowded PEG environment (an asymmetrical set-up), and this leads to the dsDNA being captured differently. This is intriguing, as it implies that the addition of PEG to the bath alone alters the local ion environment near the nanopore and the detection mechanism here that can potentially be different from previously studied models (3, 4, 15, 27). As a note to the reader, it is not the scope of this paper to elucidate the mechanism driving the signal enhancement. The mechanism leading to the signal enhancement is still elusive but it is likely influenced by an interplay of the chemical properties of PEG and its physical effects on the concentration of ionic species in the nanopore.

Here, we studied whether the nature of the electrolyte could also influence the characteristics of the translocation events. Besides KCl, LiCl is commonly used in nanopore detection of nucleic acids because of its ability to slow down their translocation speeds (26, 48). Therefore, we tested if a combination of PEG and LiCl as the supporting electrolyte could both enhance the signal-to-noise ratio and reduce the translocation speed (Figure 2A). Interestingly, we observed that LiCl did indeed reduce the translocation speed of dsDNA but with a reduced signal to noise ratio when compared to KCl (Supporting Figure 7). We also noticed that only the bath electrolyte influences the magnitude and dwell time of the single molecule events while the electrolyte within the nanopipette, where the analytes are placed, plays a negligible role (Figure 2B and C). This is important because different analytes require specific buffer conditions and ionic strength to maintain their integrity, *e.g*. DNA origami and ribosomes both requires the presence of Mg^2+^ ions to stabilise their structure (49, 50), and protein structure can become unstable under high salt concentration (51). It is also interesting to note that the increase in dwell time when LiCl is used as supporting electrolyte has been reported before, and this effect was explained by the stronger binding affinity of Li^+^ to the DNA backbone compared to other cations (26, 48). However, this scenario is unlikely to happen in our experimental setup as when dsDNA solution was diluted in presence or absence of LiCl, it produced identical population scatters in the PEG KCl and the PEG LiCl solution (Supporting Info 7), indicating that an interaction between Li^+^ and the PEG molecule in the bath is likely to be the driver of the signal enhancement. This observation has led us to hypothesise that a cooperative effect between the electrolyte and PEG in the bath solution is responsible for the signal enhancement and that the nature of bath electrolyte may affect the translocation event signals depending on their ability to interact with PEG.

**Figure 2.**
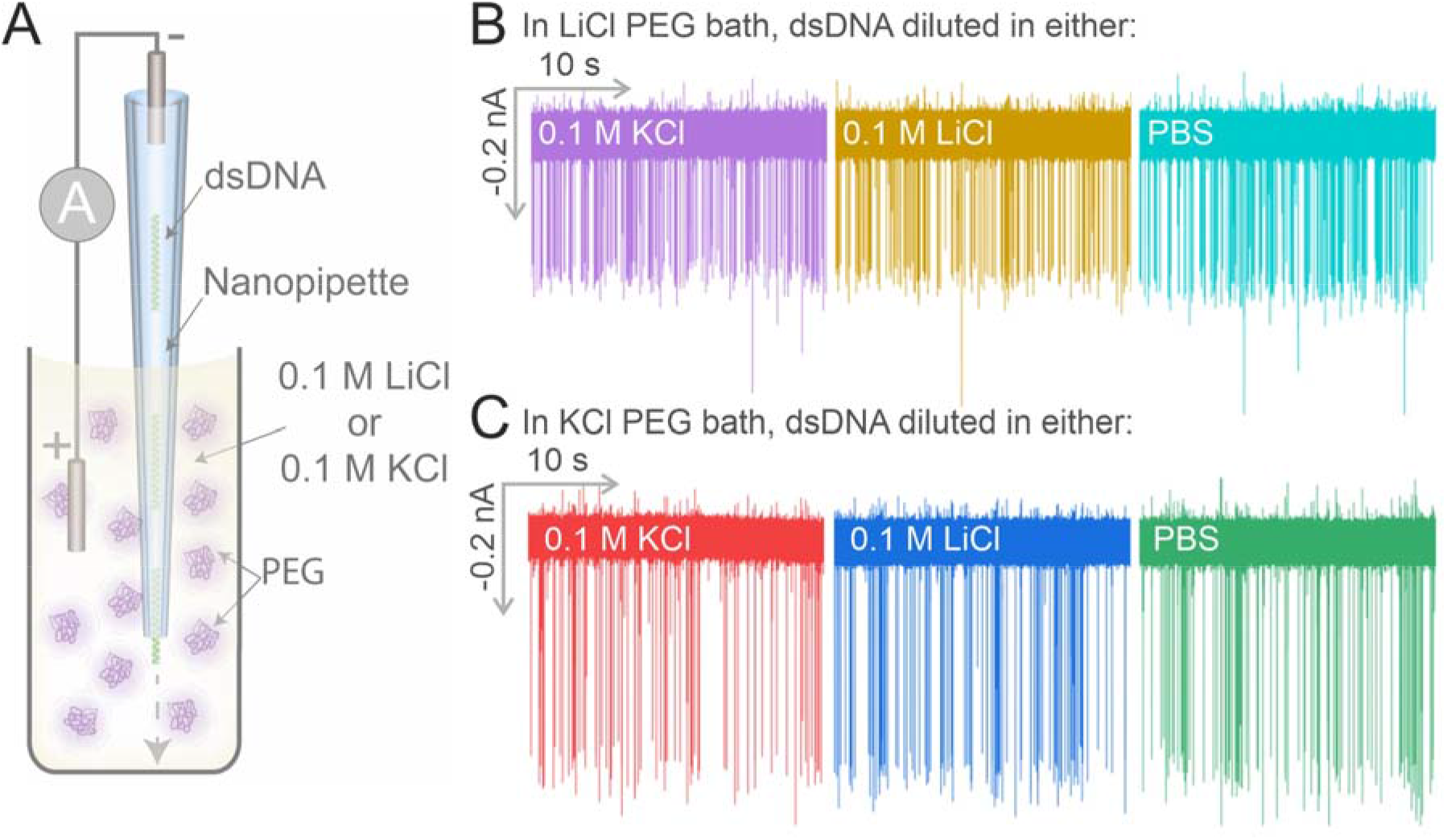
The bath electrolyte controls the characteristics of the translocation event signal. (A) The experimental set-up schematic for this experiment, the dsDNA filled nanopipette will be immersed into either a 0.1 M LiCl or a 0.1 M KCl PEG 35K bath. (B) The dsDNA was diluted in either 0.1 M KCl, 0.1 M LiCl or PBS and was immersed into a 0.1 M LiCl PEG bath to carry out translocation experiments. (C) The dsDNA was diluted in either 0.1 M KCl, 0.1 M LiCl or PBS and was immersed into a 0.1 M KCl PEG bath to carry out translocation experiments.

To test our hypothesis, we prepared a range of alkali metal halide solutions that were all dissolved at a concentration of 0.1 M in 50% (w/v) PEG 35K. All translocations were all carried out at −500 mV (Supporting Figure 8). Our results indicate that the nature of the electrolyte affects both the magnitude of the current and dwell time of the single molecule translocation events (Figure 3). We noticed that CsBr caused the largest SNR enhancement as it had the strongest amplification on the current peak maxima with the shortest dwell time, and LiCl generated the largest increase in the average dwell time with the lowest enhancement on the current magnitude. Additionally, we observe a trend where salts with higher molar masses, *e.g.* CsBr, KI, CsI, lead to stronger amplification of current peak maxima, while salts with lower masses such as LiCl, LiBr, NaF generally showed longer dwell time (Figure 3). Finally, the decrease in the dwell time and increases in the current peak maxima associates with an increase of the atomic number of the anions (*e.g*. from KF to KCl to KBr & KI, Supporting Figure 8). We also noticed that the Li^+^ and Na^+^ seem to behave differently from the K^+^ and Cs^+^ ions, based on the population scatters generated (*e.g.* LiCl & NaCl vs KCl & CsCl, Supporting Figure 8). This indicates that the molar mass of the salt, and subsequently, the size of the anions and cations could also be a contributing factor for the observed broadening of dwell time effect.

**Figure 3.**
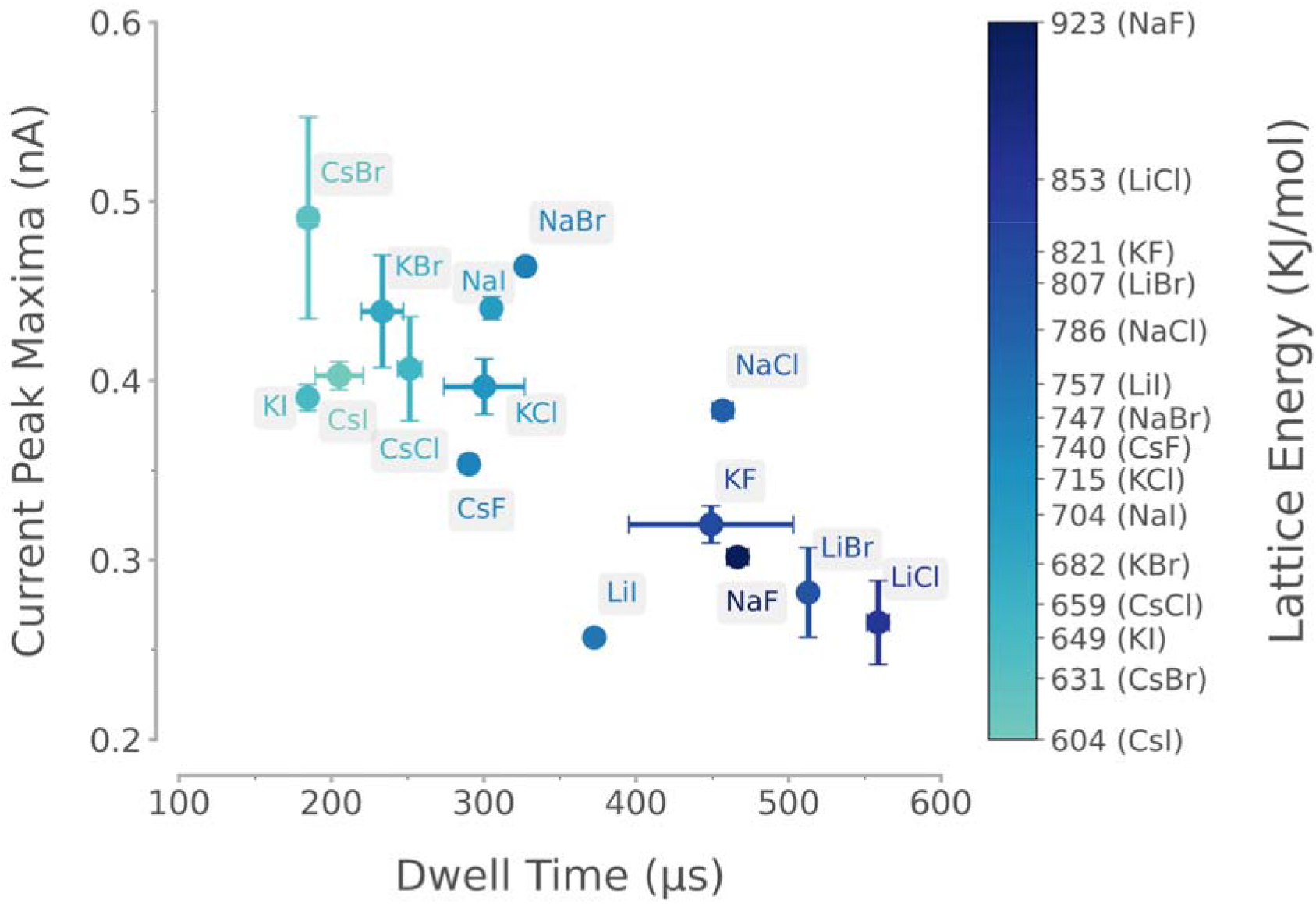
The influences of the different alkali metal halide salts on the translocation event signals. The dsDNA was diluted to 0.3 nM in 0.1 M KCl and was used to fill nanopipettes in all experiments, different 0.1 M alkali metal halides were used to generate the PEG 35K bath, these include the LiCl, LiBr, LiI, NaF, NaCl, NaBr, NaI, KF, KCl, KBr, KI, CsF, CsCl, CsBr and CsI. The KCl diluted dsDNA nanopipettes were immersed into these different salt PEG bath, and voltage of −500 mV were used to cause the dsDNA to translocate into the electrolyte bath in all cases.

The translocation of PEG through biological nanopores has been well documented (34-39) and several nanopore polymer translocation studies had pointed out to the ability of PEG to interact with cations in solution (40, 41). These studies utilised the biological nanopore α-haemolysin to detect the translocation of PEG and showed that the PEG molecules were neutral when a Li^+^ containing electrolyte was used. In contrast, PEGs were positively charged when other alkali halide metals were used, including the cations Na^+^, K^+^, Rb^+^ and Cs^+^. In 1982, Papke *et al*. studied the thermodynamic properties of the PEG-electrolyte interactions with sodium containing electrolytes and showed that the lattice energy of these electrolyte could be used to estimate their likelihood to interact with PEG (52). Lattice energy is the energy required to separate a mole of an ionic solid into gaseous ions and it is inversely proportional to the ionic compound molar mass. Based on Papke *et al*.’s observations, the electrolyte interacts with PEG if it has a lattice energy higher than 750 KJ/mol, while electrolytes with a lattice energy lower than 750 KJ/mol do not interact with PEG (43).

.All the electrolytes tested with a lattice energy above 750 kJ/mol showed high current enhancement and short dwell times, while we observed the opposite effect for electrolytes with lattice energies below 750 kJ/mol (Figure 3, Supporting Figure 9). Overall, our results show that the nature of the electrolyte is a major determinant for the dwell time and current maxima of dsDNA translocation. These differences can be associated to the interaction between the salts and PEG, as approximated by the use of lattice energy (43), indicating that a potential cooperative interaction between the electrolyte and PEG is the driver for the signal enhancement. The exact mechanism that leads to the observation is still elusive. We tested PEG of different sizes and the experiment results showed that increasing the molecular weight of the PEG associates with a stronger current enhancement, which can potentially be due to the increasing crowding effect caused by the size of the PEG (51, 52). Although PEG can be used as a macromolecular crowder, PEG is also known for its ability to chelate cations (68), a property that has been extensively studied in the field of Li-ion batteries (53), and cation chelation could potentially explain some of our observations (53, 54). Furthermore, when an electric field is applied in a solution containing PEG, the cation migration is hindered by the interactions with the polymer itself, thus causing a reduction in their mobility (53, 55, 56). To further understand the mobilities of different cations in the PEG electrolyte, Zhang *et al*. in 2015 employed a molecular dynamic (MD) simulation validated with quasi-elastic neutron scattering to simulate the mobilities of Li^+^, K^+^ and Na^+^ cations in a 50% PEG solution equivalent system (57). The study showed that cation K^+^ is significantly chelated by the PEG chain and spent nearly 5 ns interacting with the PEG chain, in contrast to Li^+^ which is a lot more mobile and only spent less than 1 ns interacting with the PEG chain (57). The observed differences in the translocation event signals between the Li^+^, Na^+^, K^+^ and Cs^+^ solutions could be due to the differences in how strongly these cations interact with the ethylene oxide group of PEG.

To conclude our work, we present two examples of novel experimentations that can be performed by taking advantage of the combination of PEG and CsBr in the bath solution.

The first key advantage of the enhanced SNR is that we could detect dsDNA in absence of a low-pass filter (bypass setting). For most solid-state nanopore translocation experiments, due to its inherent high frequency dielectric and capacitive noises (24), a low-pass filter (typically at 10KHz) must be applied to reduce the noise to allow the discrimination of the translocation events (58, 59). However, filtering comes at a cost of a reduced temporal resolution of the acquired data.

We first tested the translocation of 4.8 kbp dsDNA at −500 mV into the CsBr PEG bath with different low-pass filter setting and translocation events could still be observed even when the low-pass filter was bypassed (Figure 4A). In contrast, when we used PEG 4K in conjunction with CsBr, no translocation events could be observed without a low-pass filter (Supporting Figure 11). It is noteworthy to point out that the current noise has increased from *ca*. 0.1 nA with a 20 kHz low-pass filter to *ca*. 0.5 nA with the bypassed setting (Figure 4A). Because of the large enhancement in the SNR, we were able to increase the sampling frequency to 500 kHz, therefore enabling a 2 μs temporal resolution which is the physical limit of our amplifier and translocation events could be detected under an applied voltage as low as −300 mV (Supporting Figure 12).

**Figure 4.**
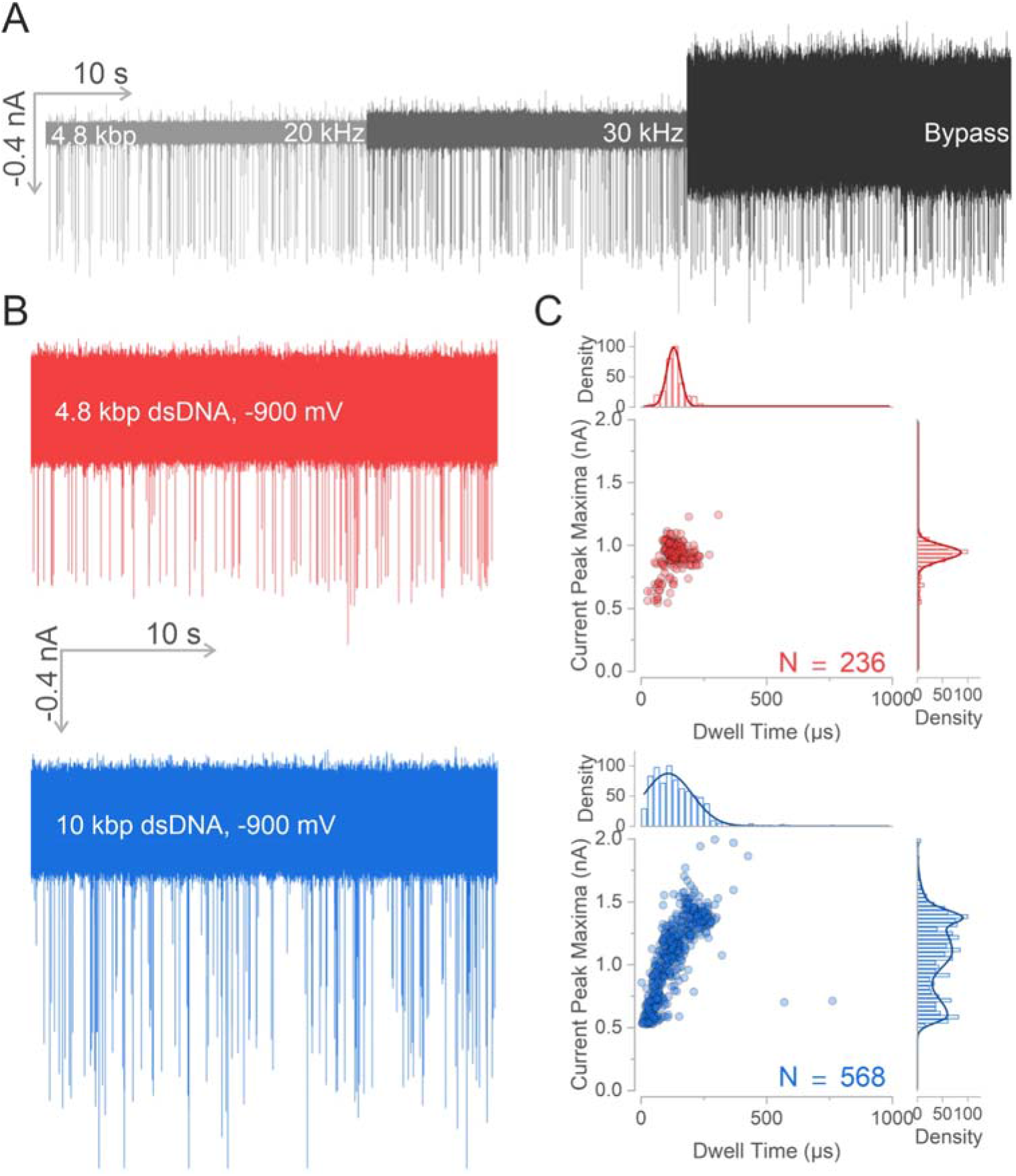
The combination of the PEG and CsBr enables the detection of dsDNA at 2 μs time resolution. (A) The translocation of 4.8 kbp dsDNA from the nanopipette into the CsBr PEG 35K bath under either 20 kHz, 30 kHz or bypass low-pass filter settings. (B) The translocation of either the 4.8 kbp dsDNA or the 10 kbp dsDNA at −900 mV into the CsBr PEG 35K bath. (C) The population scatters generated from events recorded in (B), the 4.8 kbp dsDNA generates a population centred at 0.96 nA and 130 μs. The 10 kbp dsDNA does not form a single population, instead it forms a spread ranging from 0.5 to 1.5 nA, and at least three populations can be identified at 1.4, 1.1 and 0.6 nA. All histograms were fitted with a gaussian distributions (for normally distributed data) or bigaussian distribution (for abnormally distributed data), a multi-peaks fitted distribution was generated by the convolution of multiple gaussian distribution. N is the number of events plotted.

Multiple studies have correlated the shape of the translocation event to the folding states of dsDNA (7, 8, 10-18). Since we were able to detect the translocation of 4.8 kbp dsDNA with a 2 μs temporal resolution (Figure 4A), we investigated whether the CsBr PEG bath can facilitate the detection of different dsDNA folding state. Studies showed that the translocation of dsDNA with length of *ca*. 7 kbp and above generates a multi-level event when translocated through a *ca*. 20 nm wide solid-state nanopore, and this multi-level event shape depends on the folding state of the dsDNA (12-15, 17), so we used 10 kbp dsDNA to probe the ability of our system to improve the detection of DNA folding(Supporting Figure 13).

Similar to the translocation of the 4.8 kbp dsDNA, the 10 kbp dsDNA was detectable at voltage as low as - 300 mV in the absence of the low-pass filter (Supporting Figure 14). In contrast to the 4.8 kbp dsDNA, the 10 kbp dsDNA event data does not form a single population at higher voltage of −700 mV and above (Supporting Figure 12, 14), *e.g*. three populations can be observed at −900 mV alongside the current peak maxima axis at *ca*. 1.4, 1.1 and 0.6 nA, these events spread across the current peak maxima axis ranging from 0.5 to 1.5 nA, while the 4.8 kbp dsDNA form a single population in the scatter plot centred at *ca*. 1 nA and 130 μs (Figure 4B-C). We previously reported that the addition of PEG in the bath enables the differentiation between different DNA topologies (33). The relaxed and supercoiled DNA plasmids formed two populations on the event population scatter, the separation of these two populations were due to differences in current magnitudes, here we attribute the cause of this widespread multi populations pattern in 10 kbp dsDNA to the differently folded dsDNAs during their translocations. Furthermore, the translocation events recorded at −900 mV were individually inspected and we identified different types of multi-level events (Supporting Figure 15), that are consistent with data presented in the literature linked to dsDNA folding states (7, 8, 10-18).

The second key advantage of the increased SNR caused is that we could detect dsDNA as short as 75bp. The detection of short dsDNA (<500 bp) is challenging with nanopipettes because the SNR is often very small (60) and the translocation of short DNAs through the nanopore is too fast to be detected (28, 29, 61) even with state-of-the-art electronics (62).

Here, we first generated a short dsDNA strand of 500 bp (Supporting Figure 16) and diluted it to a concentration of 0.3 nM in 0.1 M KCl. Beside the CsBr PEG bath, the KCl and LiCl PEG baths were also tested for the translocation of the 500 bp dsDNA for comparison, we anticipated that CsBr would yield the highest molecule count as it enhanced the current magnitude the most among all the tested electrolytes (Figure 3). The same nanopipette filled with the 500 bp dsDNA was used for translocation experiments in all three salt PEG baths filtered with a 20 kHz low-pass filter. The detection of the 500 bp dsDNA was successful when a voltage of −500 mV was used in all baths and the three bath translocation events all formed a single population at *ca*. 100 μs and 0.1 nA (Figure 5A-B). Although the 500 bp dsDNA translocation events were all detectable, the molecule count in CsBr was *ca*. 7-fold higher than either KCl or LiCl (Supporting Figure 17).

**Figure 5.**
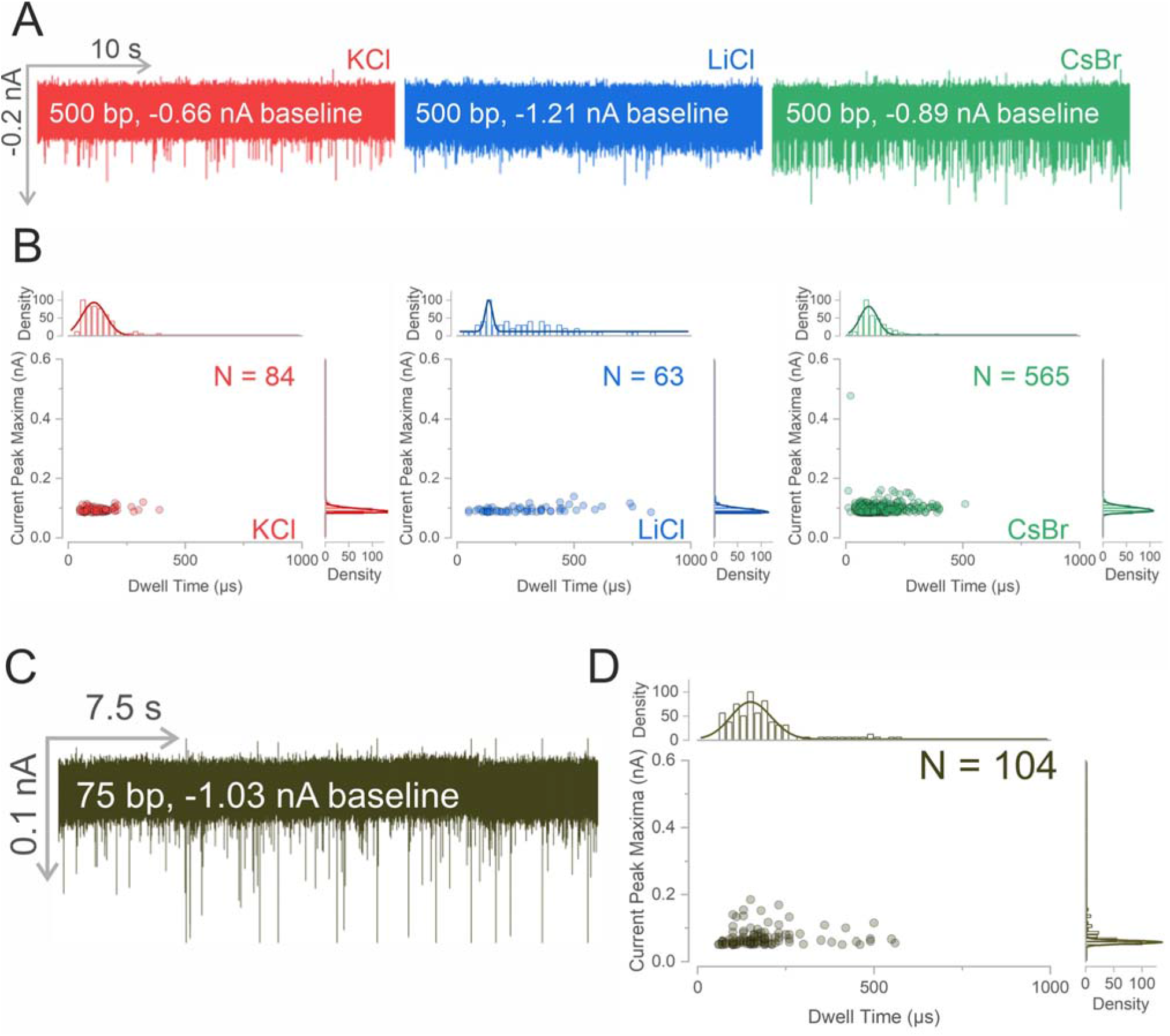
CsBr improves the sensitivity of the nanopipette for the detection of short dsDNA. (A) The 500 bp dsDNA was diluted in 0.1 M KCl and immersed into either the 0.1 M KCl, 0.1 M LiCl or 0.1 M CsBr PEG solution under a −500mV applied voltage (B) The population scatter for the translocation events, fewer than 100 events were recorded for the KCl and LiCl generated PEG 35K solution. (C) The translocation of 75 bp dsDNA, the 75 bp dsDNA was diluted in 0.1 M KCl to 1 nM and used to fill the nanopipette, the translocation of the dsDNA was carried out inside a 0.1 M CsBr PEG 35K solution by applying a voltage of −700 mV (D) The population scatter of the translocation of the 75 bp dsDNA, a total of 104 events were recorded in 30 seconds. All histograms were fitted with a gaussian distributions except D in which the dwell time histogram was fitted with bigaussian distribution. N is the number of events plotted.

The 75 bp dsDNA was diluted to 0.3 nM in 0.1 M KCl and the nanopipette was immersed into the CsBr PEG bath. Translocation events were detected when applied a voltage of −500 mV (Supporting Figure 18). Under the applied voltage of −700 mV, the events formed a population on the scatter with an average peak maximum centred at *ca*. 0.05 nA and a dwell time of about 200 μs (Figure 5D). Our results showed that CsBr PEG bath enhanced SNR in the current magnitude of the 75 bp dsDNA, but also sufficiently reduced the translocation time to become above 66 μs. The detection of short dsDNA ranging from 100 to 500 bp have been recently demonstrated with hydrogel modified nanopipettes (28, 29), here we show that dsDNA as short as 75 bp can also be detected efficiently by an unmodified 25 nm diameter nanopore by simply using CsBr as supporting electrolyte in 50% PEG 35K.

## CONCLUSION

We have demonstrated that the chemical properties of the electrolyte in the presence of 50% PEG35K significantly modify the translocation dynamics of dsDNA, as characterized by the translocation event current peak magnitude and the dwell time. Our results indicate that a cooperative effect between the electrolyte and PEG is responsible for driving the signal enhancement in a solid-state nanopore. We observed that the translocation dynamics can be explained after considering the cation binding properties of PEG, and that that the strength of the interaction between a cation and PEG can be used as a predictor for the observed signal enhancement. We further demonstrated that using CsBr as the supporting electrolyte in a PEG 35K bath enables the detection of 10 kbp dsDNA at 2 μs temporal resolution and the detection of dsDNA as short as 75 bp. Our approach relies on commercially available electronics and with a solid-state nanopore of 25 nm in diameter that can be easily and reproducibly fabricated at the bench and takes advantage of the careful selection of the bath conditions in terms of molecular weight of the PEG and nature of the electrolyte. Therefore, our approach can be easily adopted in laboratories across the world and potentially be integrated with advanced custom-built electronics that allows sampling at MHz frequencies (11). We envision that this knowledge should facilitate the study of sub-molecular translocation dynamics of single molecules through nanopores and the analysis of short nucleic acids for clinical applications.

## METHODS

The supporting information contains detailed methods on: the generation of the PEG solutions, the preparation of the dsDNA, ionic current traces and scanning electron microscopy micrographs of the nanopipette used.

### Nanopipette fabrication

Quartz capillaries of 1.0 mm outer diameter and 0.5 mm inner diameter (QF100-50-7.5; Sutter Instrument) were used to fabricate the nanopipette using the SU-P2000 laser puller (World Precision Instruments). A two-line protocol was used, line 1: HEAT 750/FIL 4/VEL 30/DEL 150/PUL 80, followed by line 2: HEAT 625/FIL 3/VEL 40/DEL 135/PUL 150. The pulling protocol is instrument specific and there is variation between different SU-P2000 pullers.

### Ion current trace recording and analysis

The translocation experiment follows similar procedure from the previous publication (33). The nanopipettes were all filled with 0.3nM dsDNA diluted in either 0.1 M KCl (P/4240/60; Fisher Scientific), 0.1 M LiCl (CHE2360; Scientific Laboratory Supplies) or 1XPBS (D8537; Sigma Aldrich) and fitted with a Ag/AgCl working electrode. The nanopipettes were immersed into the electrolyte bath with a Ag/AgCl reference electrode. The ionic current trace was recorded using a MultiClamp 700B patch-clamp amplifier (Molecular Devices) in voltage-clamp mode. The signal was filtered using low-pass filter at either 30 kHz, 20 kHz, 10 kHz or by-pass setting and digitized with a Digidata 1550B at a 100 kHz (10 μs bandwidth) or 500 kHz (2 μs bandwidth) sampling rate and recorded using the software pClamp 10 (Molecular Devices). The current trace was analysed using a custom written MATLAB script provided by Prof Joshua B. Edel (Imperial College London). For translocation events analysis, the threshold level was defined at least 5 sigma away from the baseline, only events that were above this threshold would be identified as the translocation of the molecule.

## ASSOCIATED CONTENT

### Supporting Information

All ionic current trace data can be found in the University of Leeds data repository. Supporting Information (Detailed Methods and Supporting Figures) (PDF)

## AUTHOR INFORMATION

### Corresponding Author

Eric W. Hewitt

Paolo Actis

### Author Contribution

C.C.C. designed and performed all experiments and data analysis. F.M., D.S. and M.A.E. helped with data analysis. P.A., E.H.W, S.E.R. designed the experiments and supervised the research. All authors wrote and corrected the manuscript.

## Acknowledgements

We thank Prof Joshua B. Edel (Imperial College London) for generously providing the MATLAB script used for event analysis in this study. We thank Dr Nataricha Phisarnchananan (University of Leeds) for performing the viscosity measurement of the electrolyte. We thank the members of Radford, Hewitt and Actis group for helpful discussions.

## ABBREVIATIONS

PBS: Phosphate Buffered Saline
PEG: Poly(ethylene) glycol
dsDNA: double stranded DNA
SNR: signal-to-noise ratio
MW: molecular weight

