## Supporting Information for "Cooperative electrolyte-PEG interactions drive the signal amplification in a solid-state nanopore"

### Table of Content

### Supporting Figures

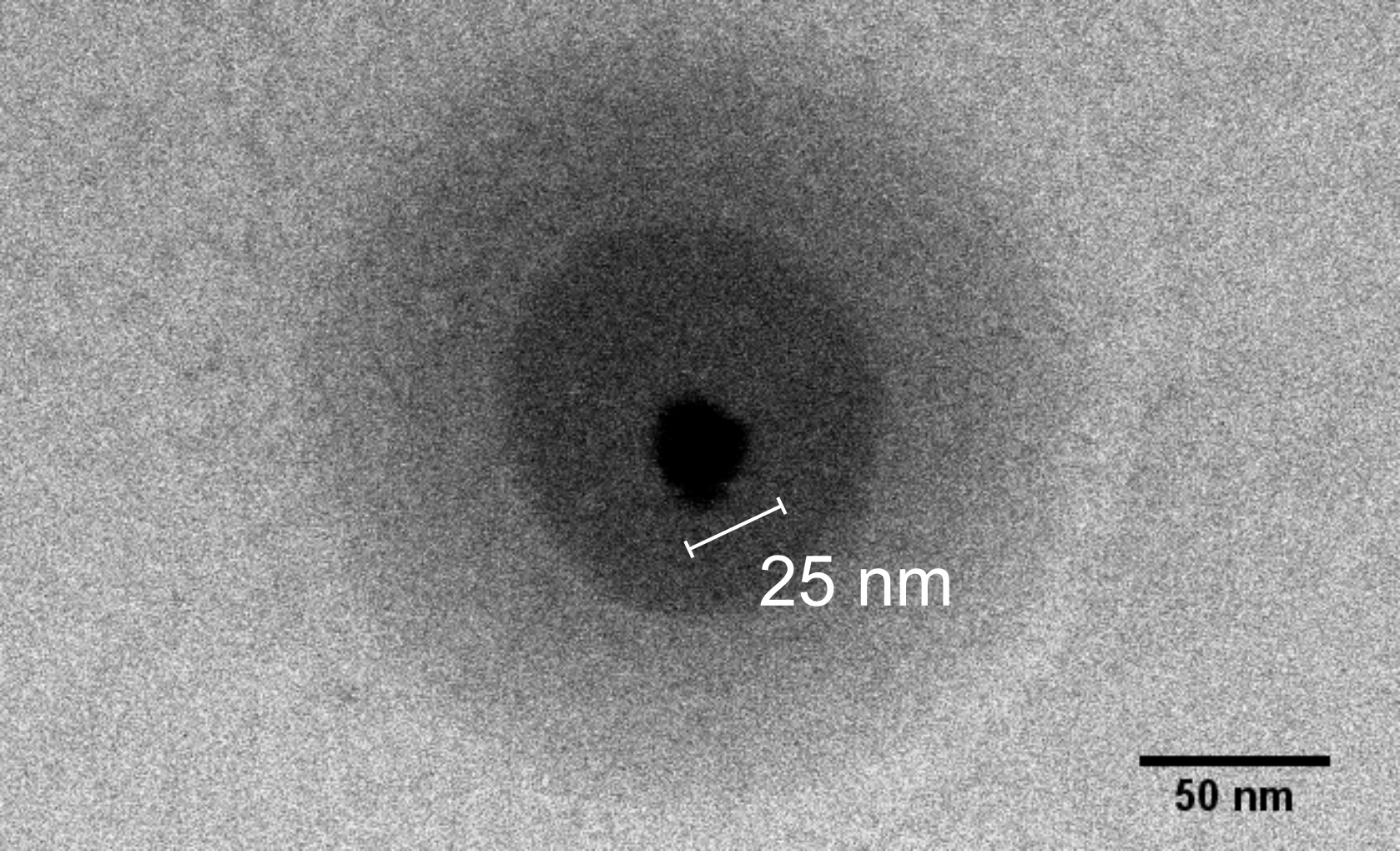

Supporting Figure 1. Scanning electron microscopy image of a nanopipette used in this study. The nanopipette used in this study was approximately 25 nm in diameter.

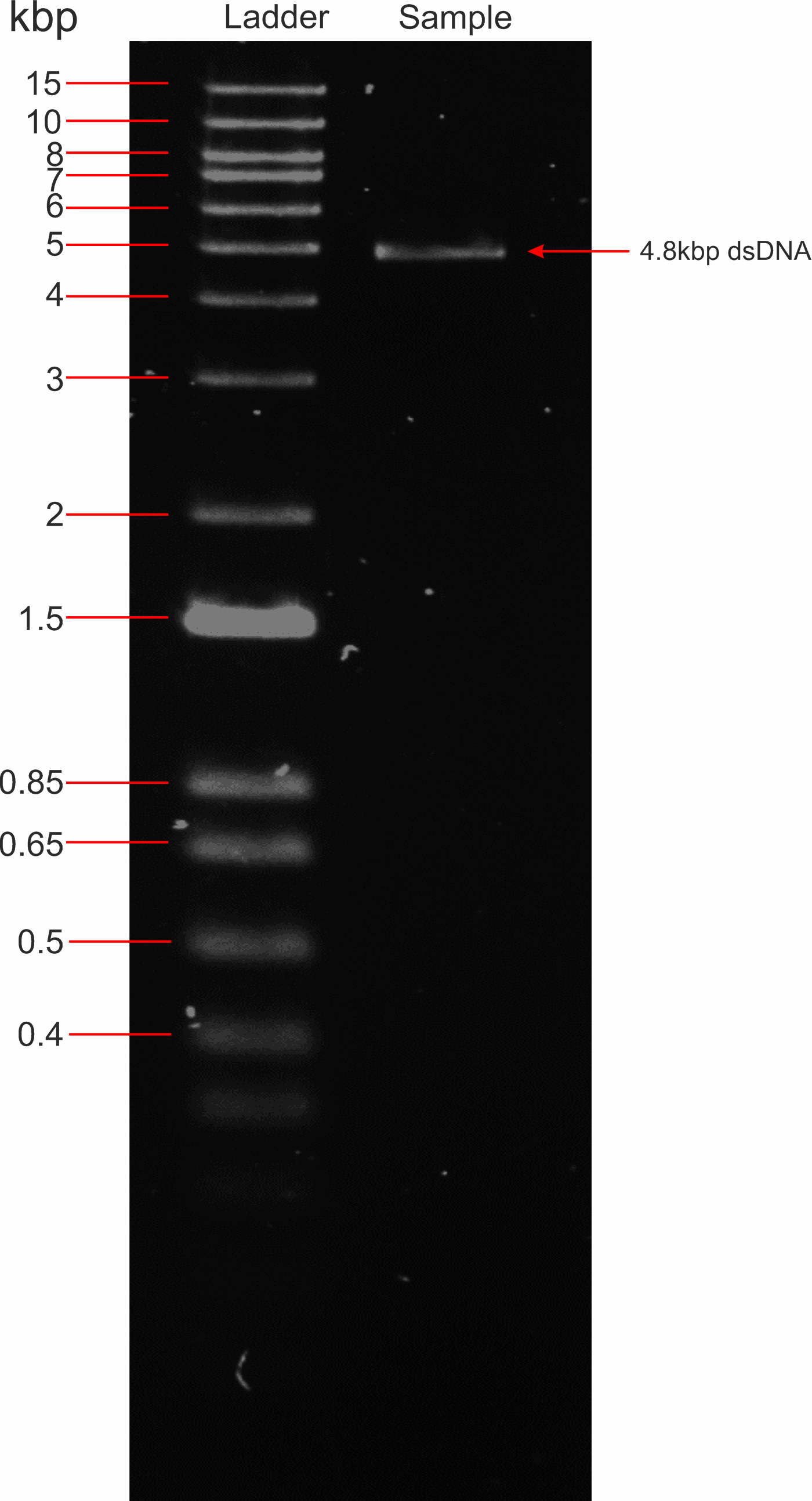

Supporting Figure 2. Gel electrophoresis analysis of the linearized pmCherry-NLS. 1 µg of the linearized plasmid (following the restriction digestion using the enzyme BamHI-HF) was analysed by gel electrophoresis. A DNA ladder was run alongside the digested plasmid sample. The digested plasmid shows a single band at around 5 kbp, which corresponds to the linearized pmCherry-NLS plasmid of *c.a.* 4.8 kbp (4.755 kbp).

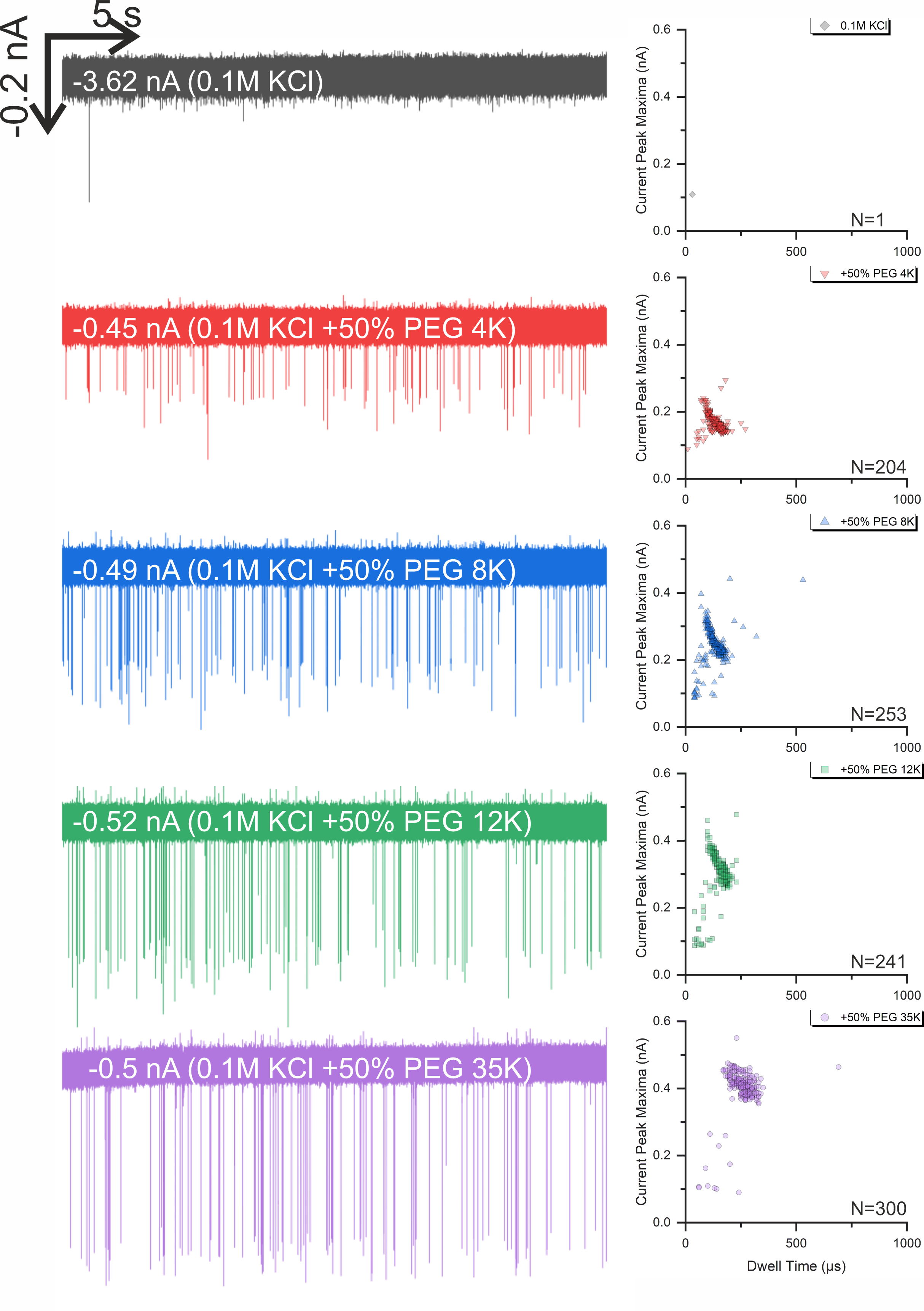

Supporting Figure 3. Translocation of the 4.8 kbp dsDNA in 0.1M KCl and the PEG crowded 0.1M KCl bath. 0.3 nM 4.8 kbp dsDNA diluted in 0.1 M KCl was used to fill the nanopipette and then the nanopipette was dipped into either the 0.1 M KCl bath (grey), +50% (w/v) PEG 4K 0.1 M KCl bath (red), +50% (w/v) PEG 8K 0.1 M KCl bath (blue), +50% (w/v) PEG 12K 0.1 M KCl bath (green), or +50% (w/v) PEG 35K 0.1 M KCl bath (purple). A -500 mV was used to cause the dsDNA to translocate from the inside of the nanopipette to the electrolyte bath. A 30 seconds trace is shown for each trace alongside with its associated population scatter. The number insert indicates the baseline current level. N indicates the number of events recorded.

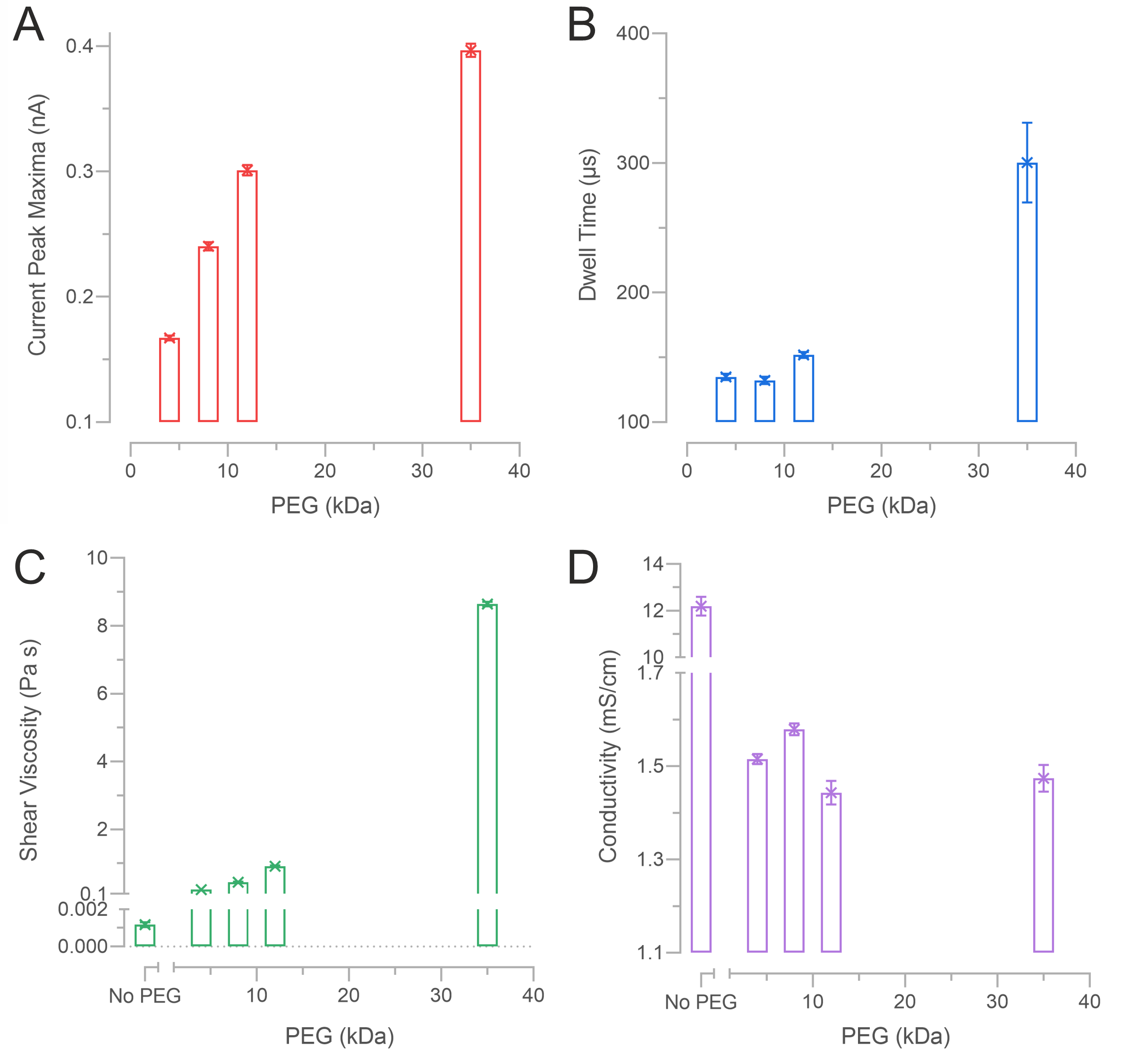

Supporting Figure 4. Summarised dsDNA translocation characteristics and physical properties of the different PEG sizes crowded 0.1M KCl bath. The average current peak maxima (A) and the dwell time (B) of the translocation events of the 4.8 kbp dsDNA carried out in the 0.1M KCl bath crowded with different sizes of PEG at 50% (w/v). The shear viscosities (C) and the conductivities (D) of the 50% (w/v) PEG crowded 0.1M KCl baths. Error bars indicate the S.E.M.

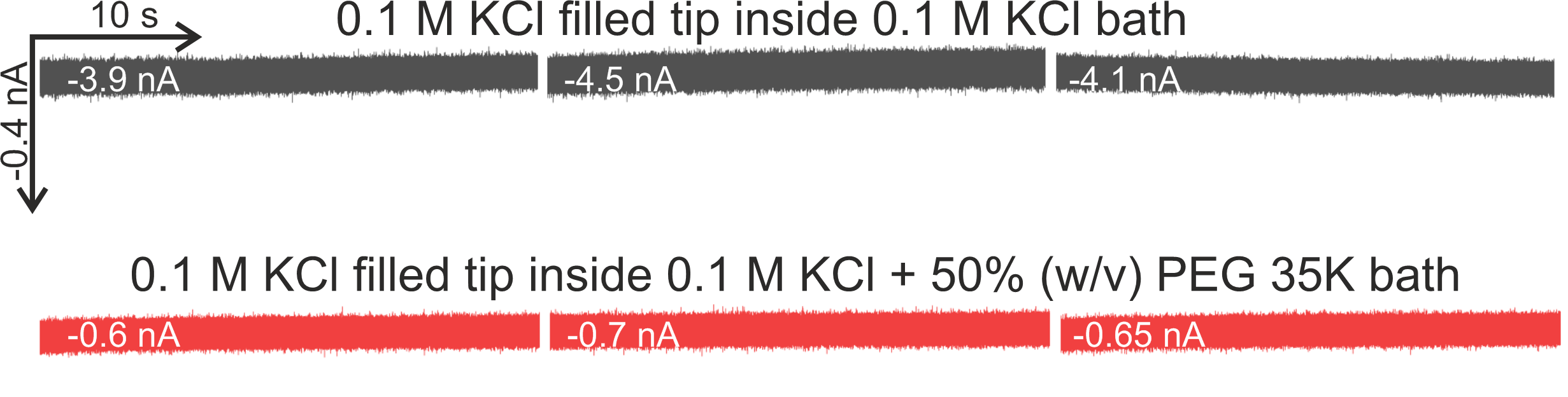

Supporting Figure 5. 0.1M KCl filled tip translocation control testing in either the 0.1M KCl or the PEG 35K crowded bath. 0.1M KCl was used to fill the nanopipette and then dipped into either the 0.1M KCl or the PEG 35K crowded bath, -500 mV was applied for 30 seconds per trace. No translocation events was recorded. Three 30 seconds traces are shown for each condition.

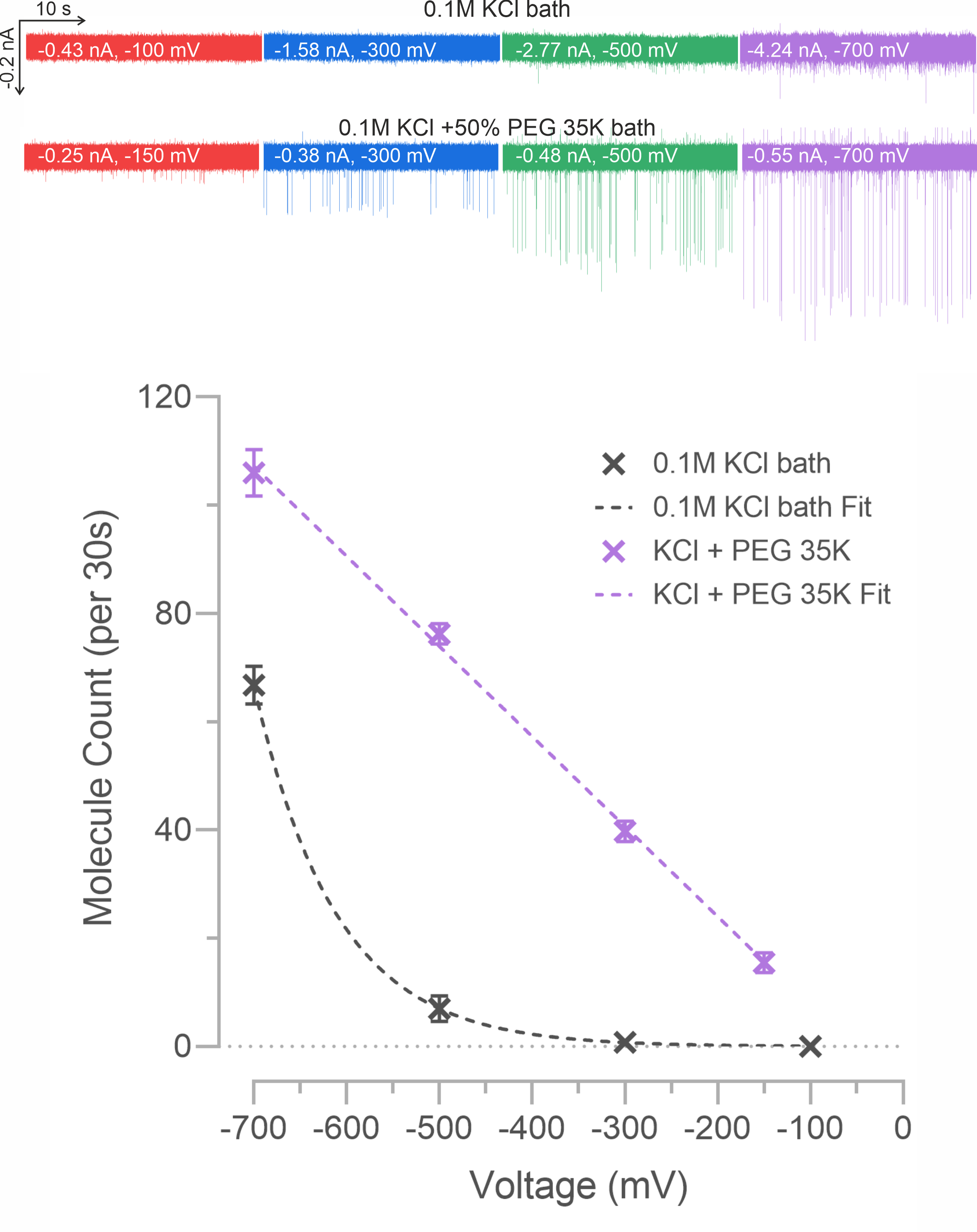

Supporting Figure 6. The voltage dependent translocation of the 4.8 kbp dsDNA with the nanopipette. 0.3nM dsDNA filled nanopipette was dipped into either the 0.1 M KCl or the 0.1M KCl +50% (w/v) PEG 35K bath. Four different magnitudes of voltage ranging from -100 to -700 mV were used to cause the dsDNA to translocate from the nanopipette to the bath. A 30 seconds trace is shown for each voltage, the number insert indicates the baseline current level.

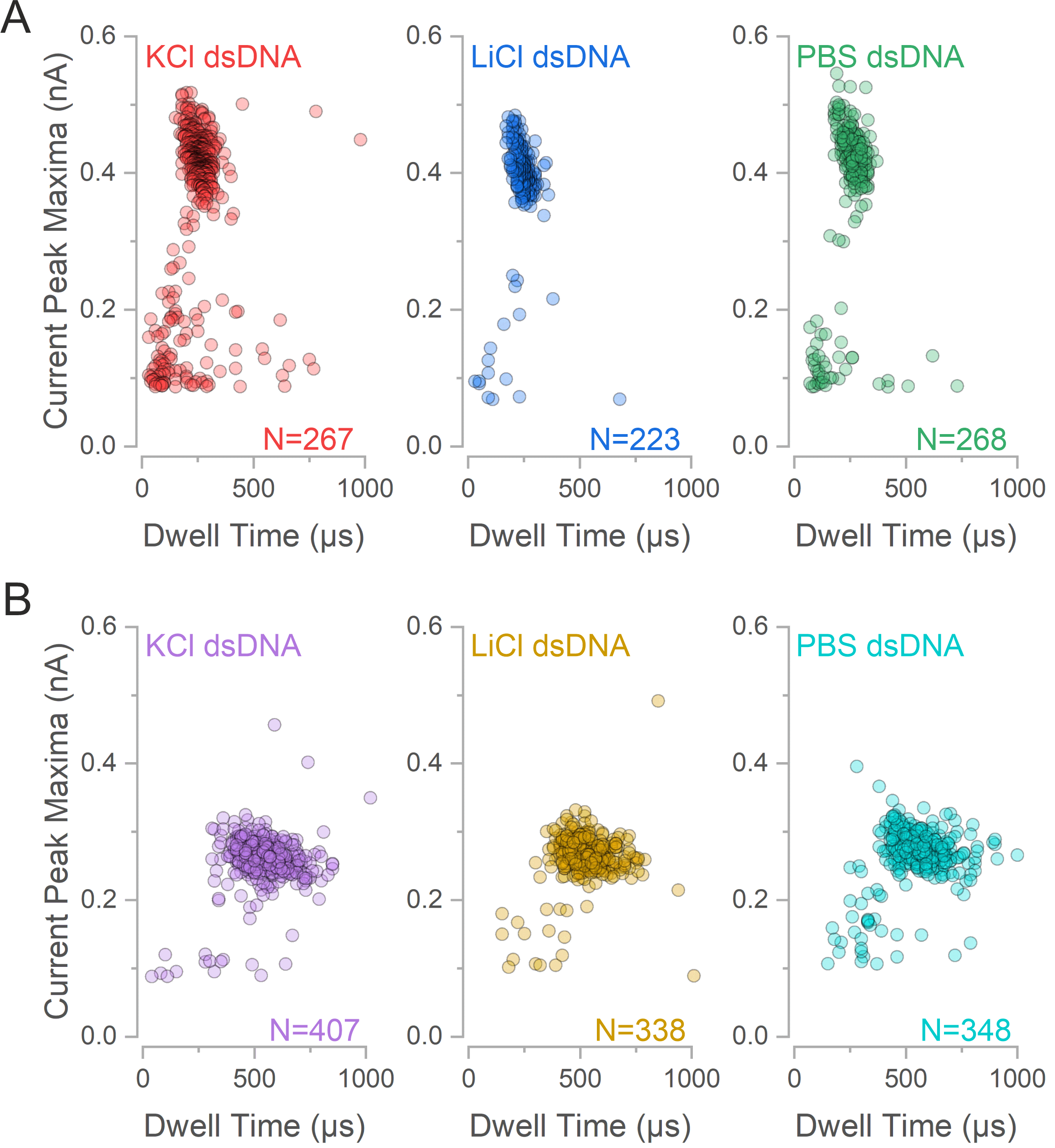

Supporting Figure 7. Population scatter plots of the translocation of differently diluted dsDNA into the KCl or LiCl PEG 35K crowded bath. The dsDNA was diluted in three different electrolyte: 0.1M KCl, 0.1M LiCl or the 1X PBS to 0.3 nM. Then the dsDNA was used to fill the nanopipette and dipped into either a 0.1M KCl +50% (w/v) PEG 35K (A) or 0.1M LiCl +50% (w/v) PEG 35K (B) to carry out dsDNA translocation. -500 mV was applied in all cases. N indicates the number of translocation events recorded.

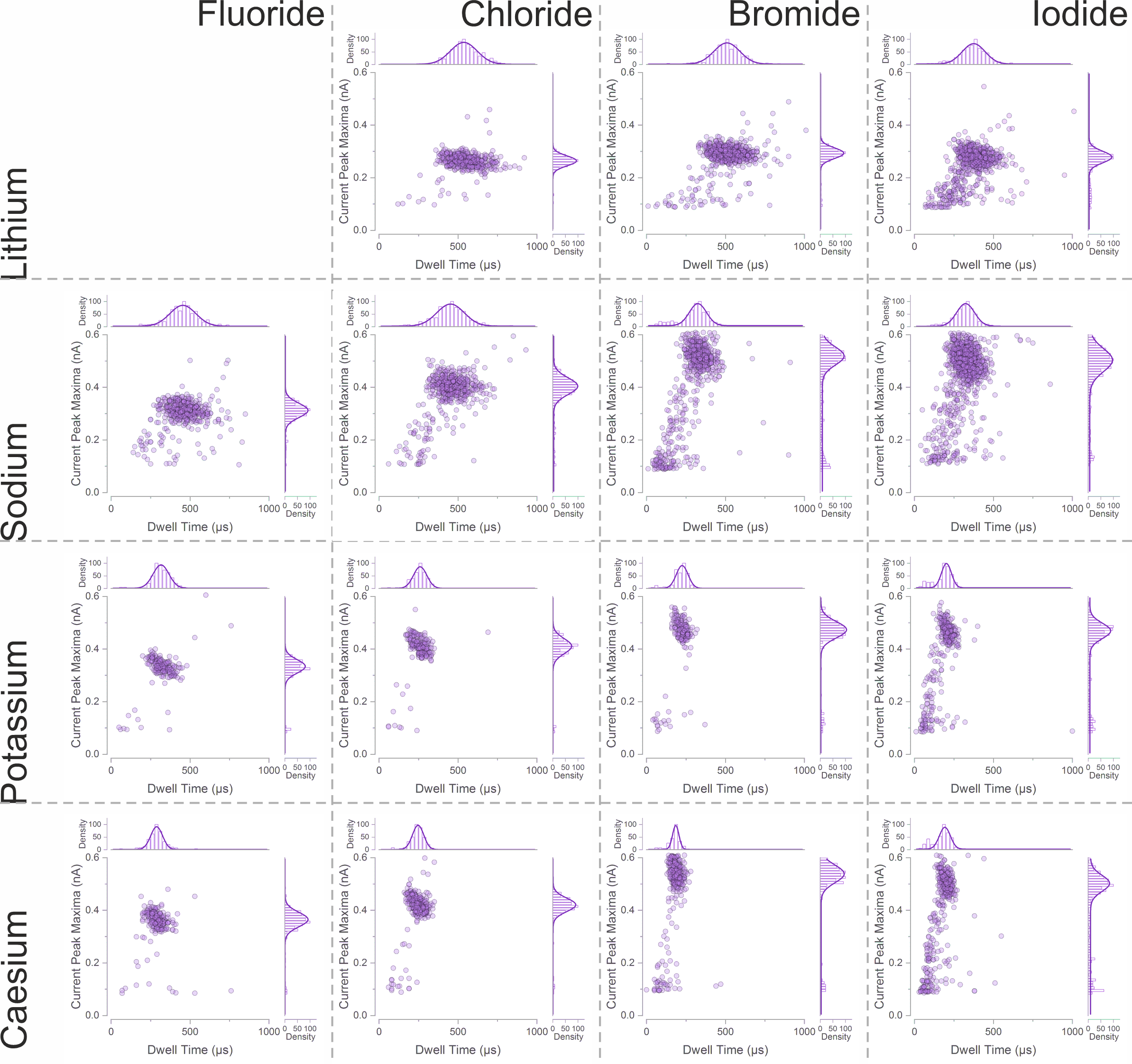

Supporting Figure 8. The event population scatter for all the tested alkali halide combination under the PEG 35K crowded condition. All nanopipettes used here were filled with 0.3 nM dsDNA diluted in 0.1M KClm, and the translocation of the dsDNA was carried out by applying a voltage of -500 mV to cause the dsDNA to migrate from the nanopipette to the bath. The alkali metals and halides species in the indicates the salt combination used to generate the 0.1M alkali metal halides +50% (w/v) PEG 35K crowded bath. The graphs are organised such that the halide atomic radii increase from left to right and the alkali metal atomic radii increase from top to bottom.

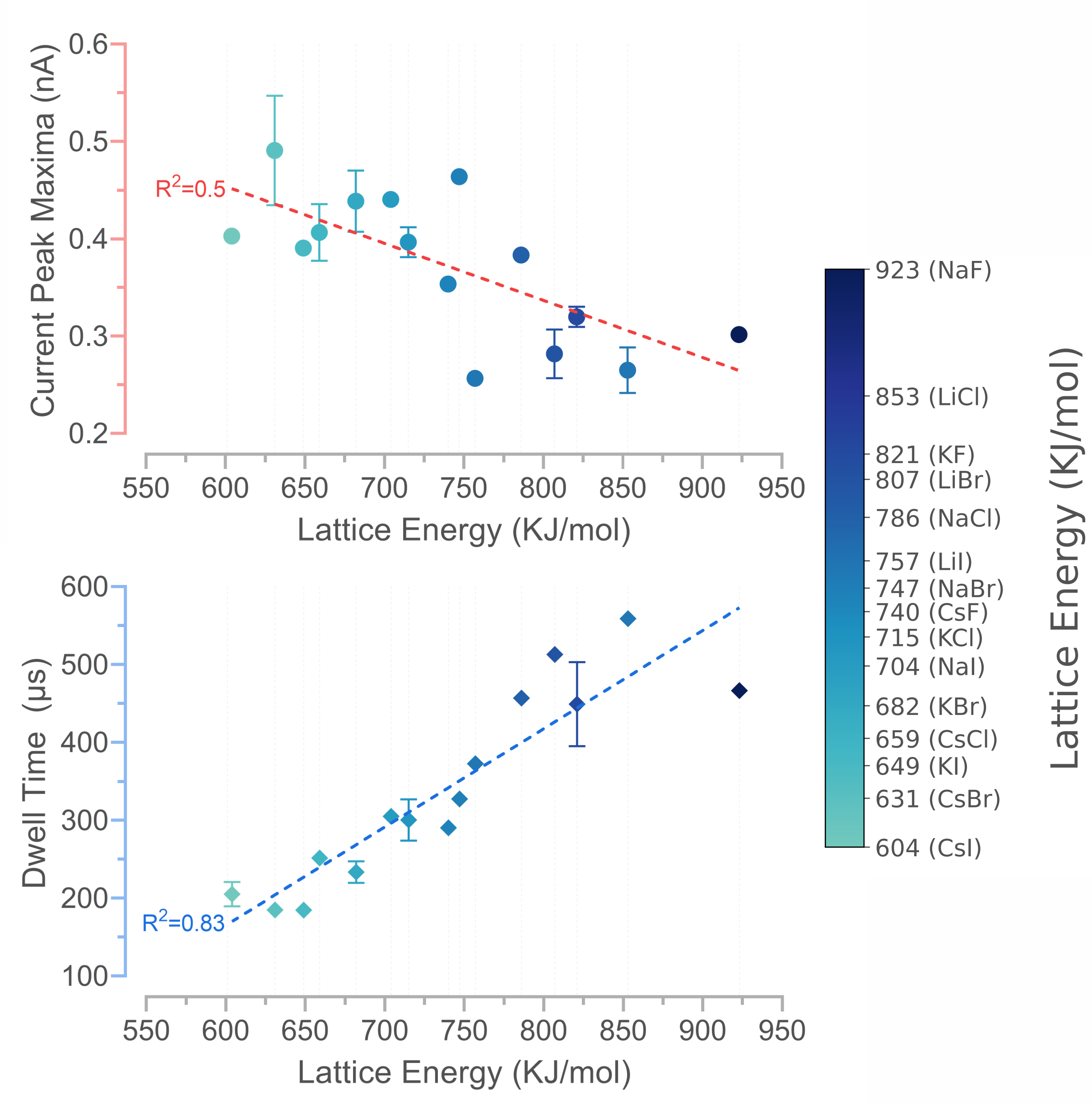

Supporting Figure 9. Amplitude and dwell time as a function of the lattice energy of the supporting electrolyte

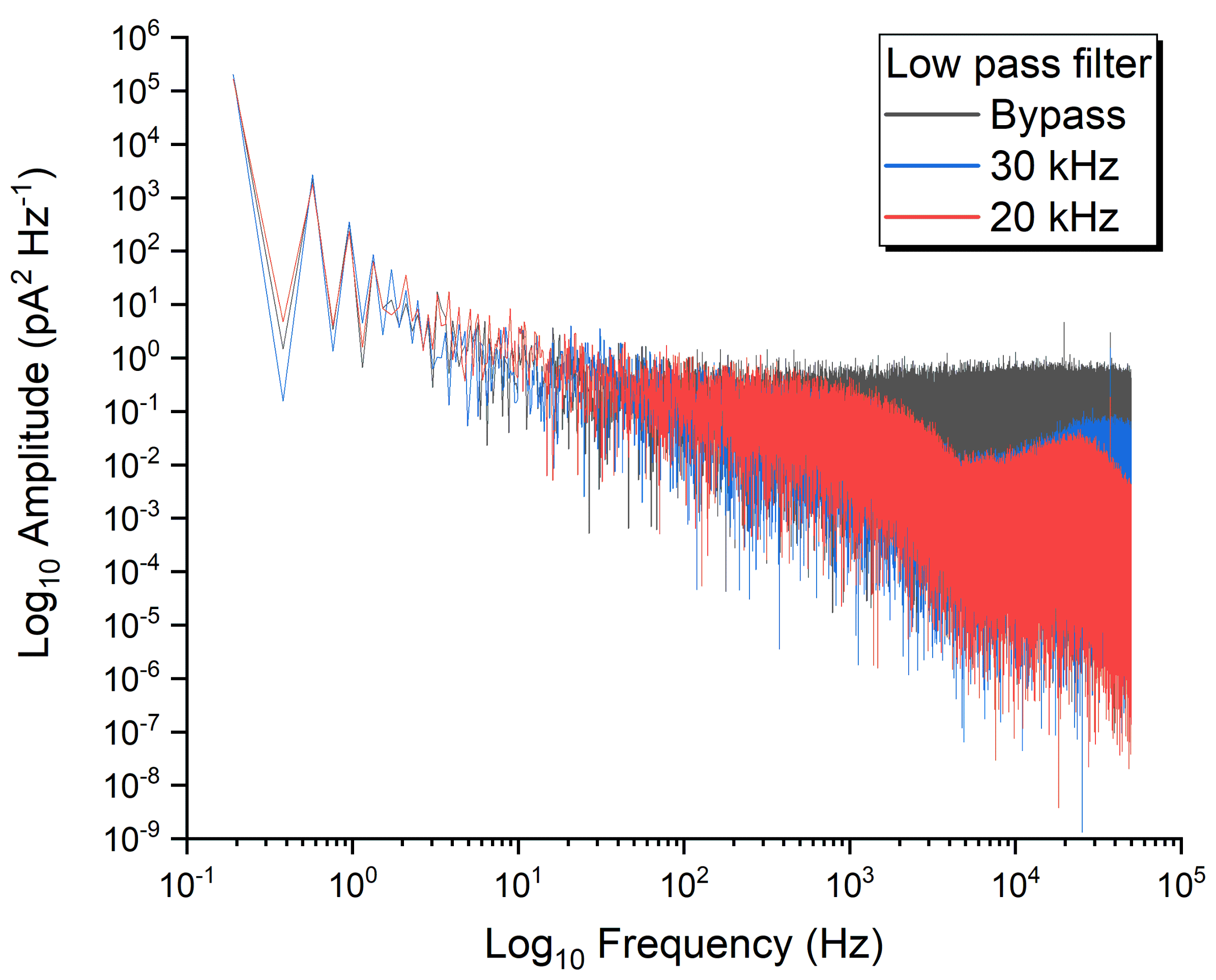

Supporting Figure 10. nanopore noise power spectral density for the low-pass filter employed in this study

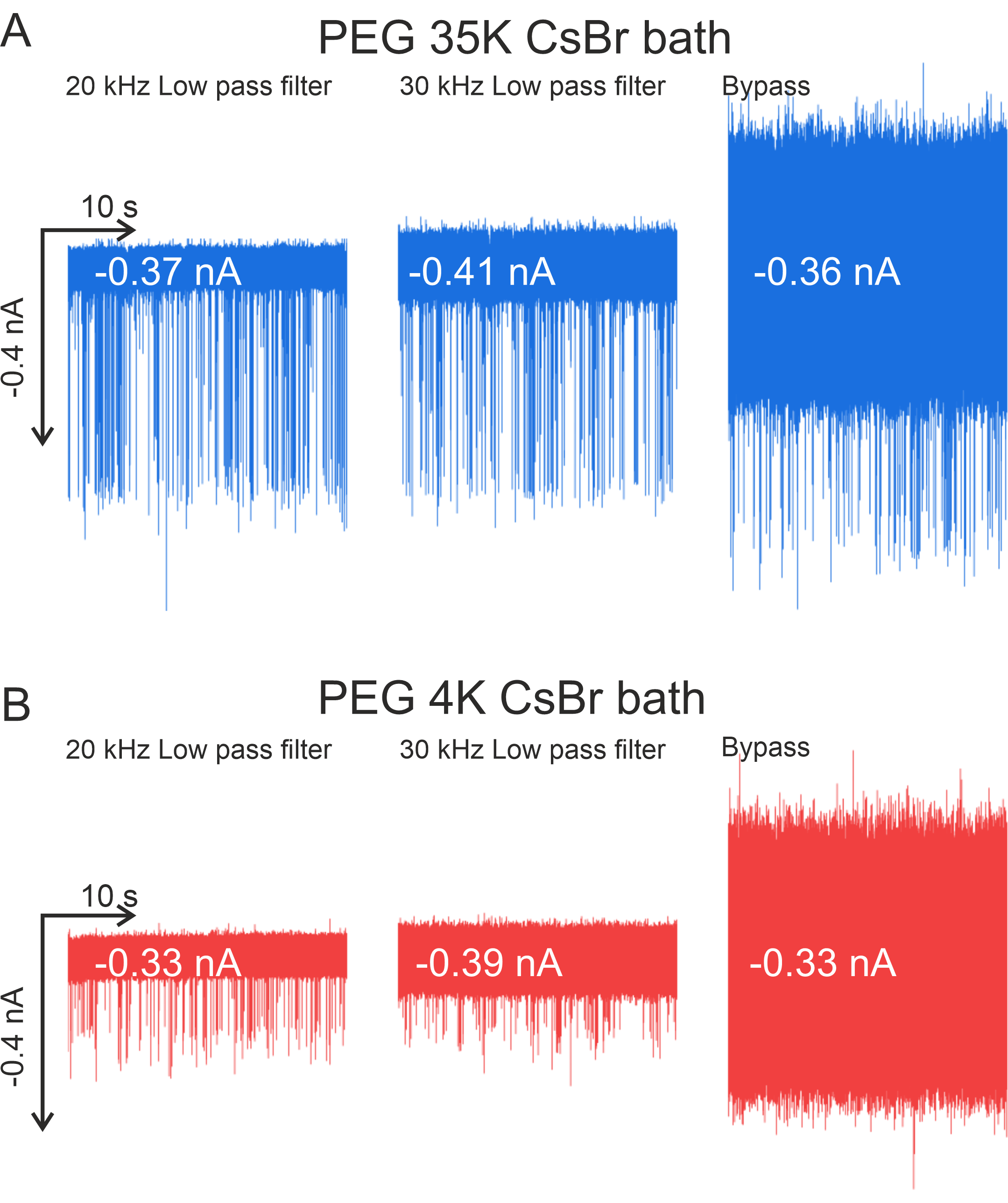

Supporting Figure 11. The translocation of the 4.8 kbp dsDNA into the CsBr PEG bath under different low-pass filter settings. (A) The translocation of the dsDNA at -500 mV into a PEG 35K crowded CsBr bath with either 20 or 30 kHz or without the low-pass filter. (B) The translocation of the dsDNA into the PEG 4K crowded CsBr bath with either 20 or 30 kHz or without the low-pass filter.

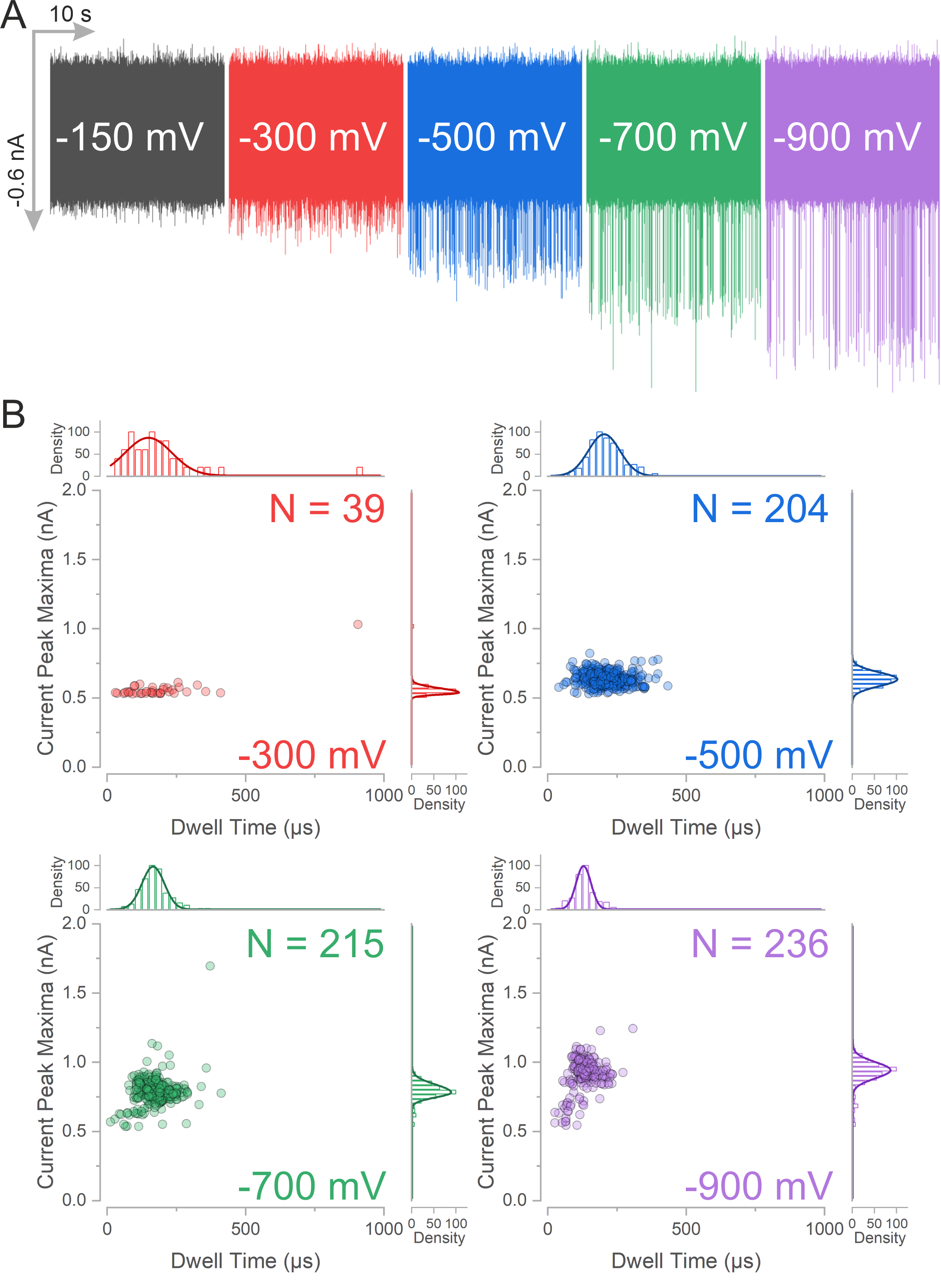

Supporting Figure 12. The voltage dependent translocation of the 4.8 kbp dsDNA into the PEG crowded CsBr bath. (A) Five different voltages from -150 to -900 mV to trigger the translocation of the dsDNA. (B) The population scatters of the translocation of the dsDNA.

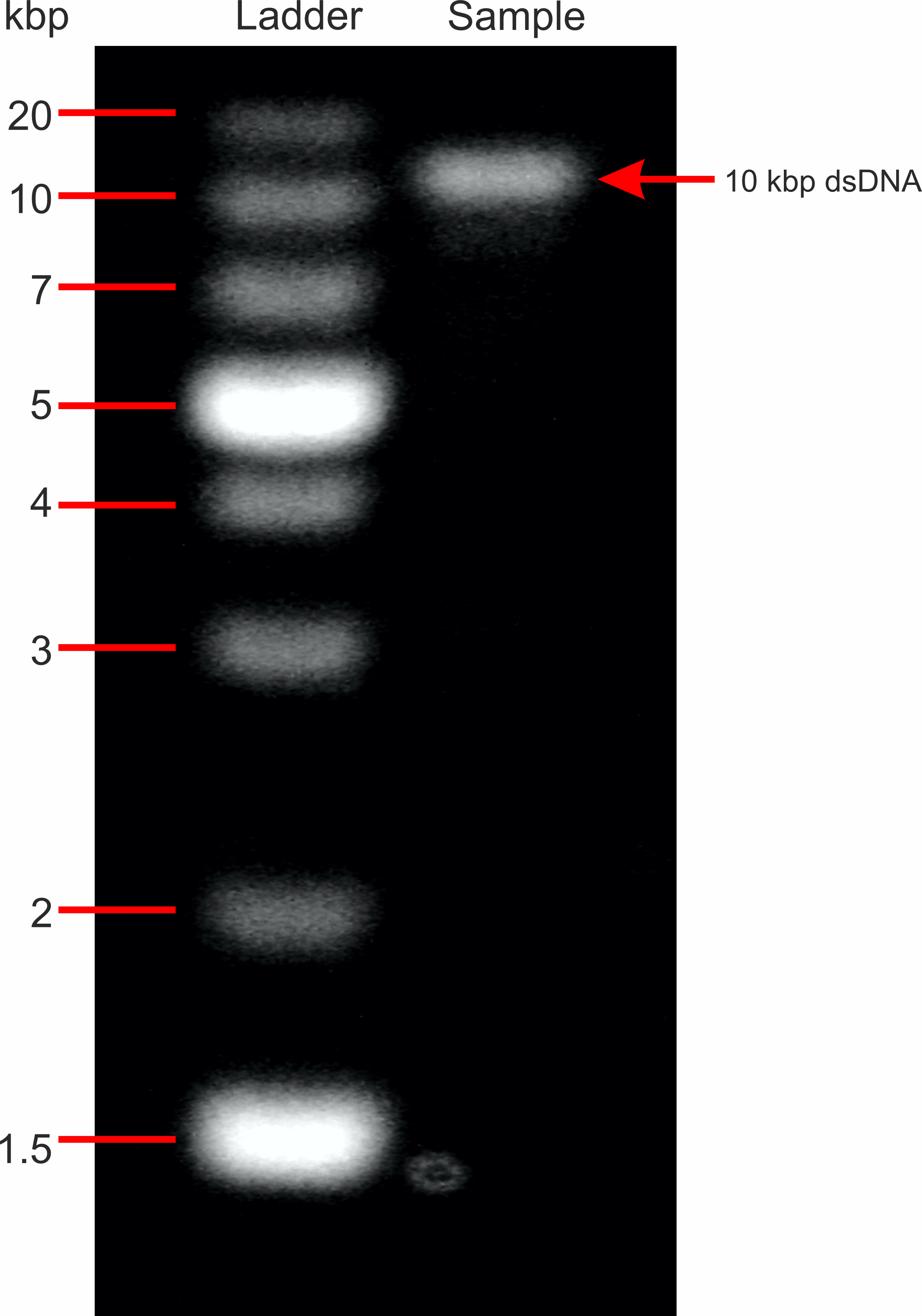

Supporting Figure 13. Gel electrophoresis analysis of the gel extracted 10 kbp dsDNA. 1 µg of the gel extracted 10 kbp dsDNA was analysed by gel electrophoresis. A DNA ladder was run alongside. The extracted sample migrated at around 10 kbp.

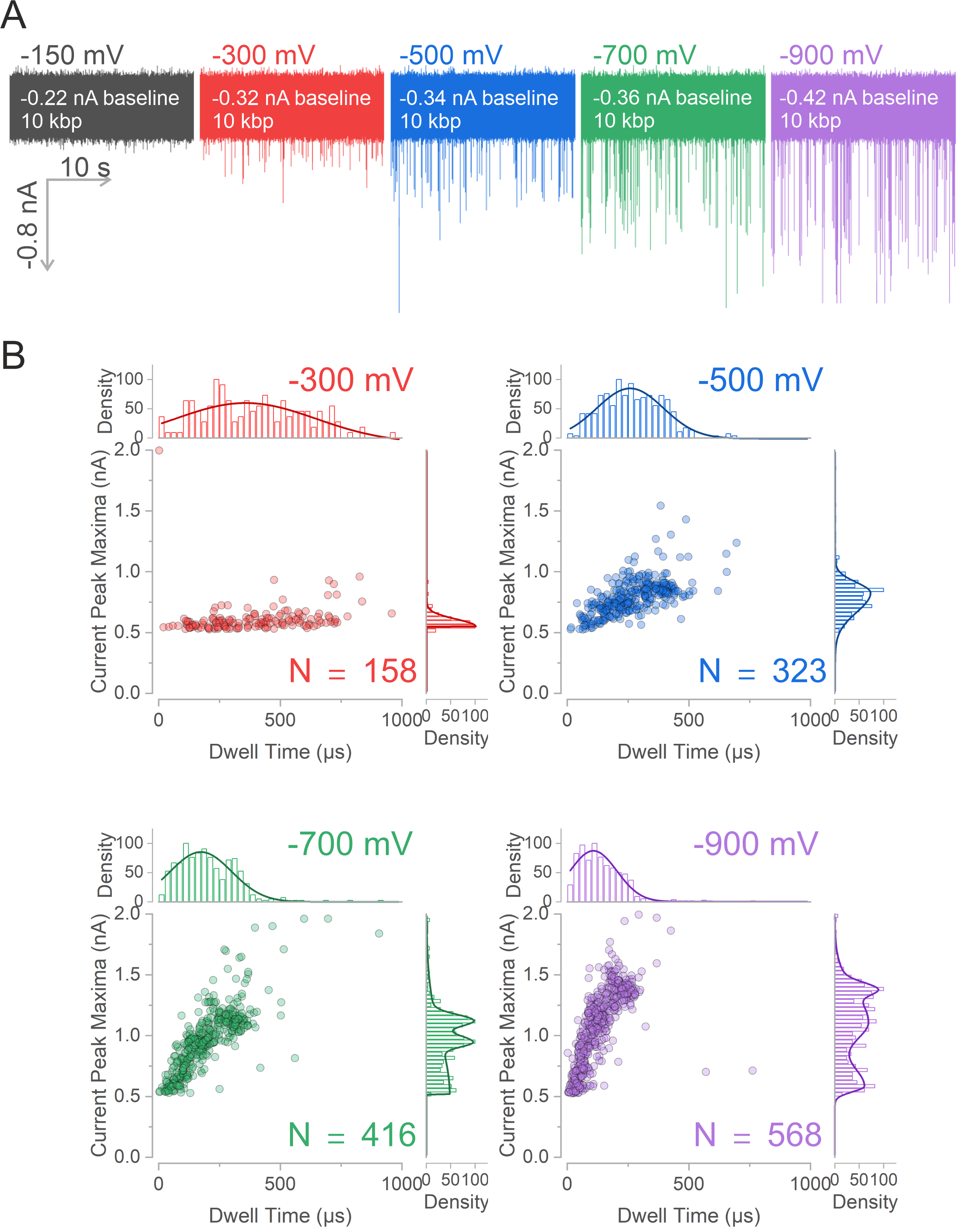

Supporting Figure 14

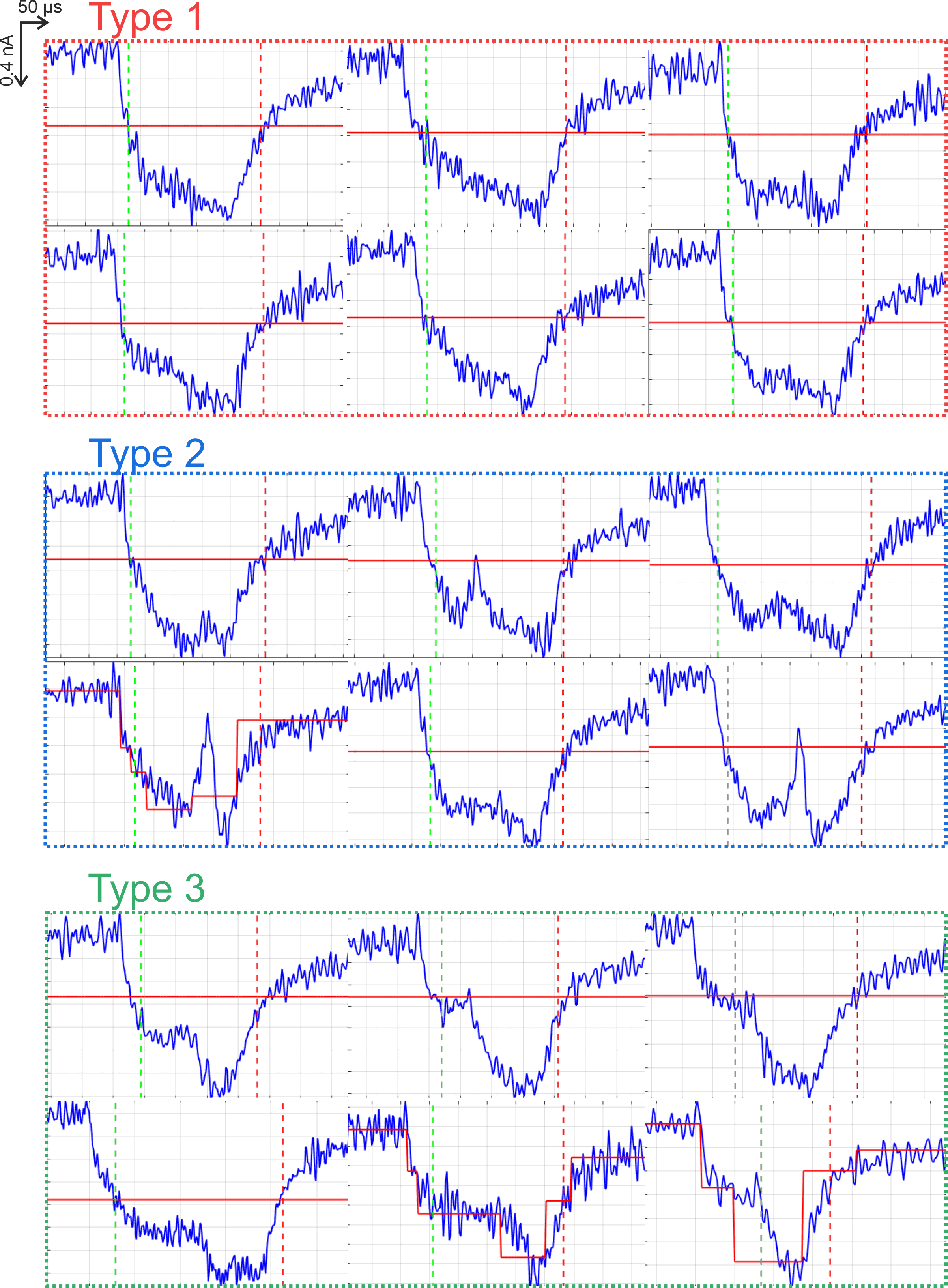

Supporting Figure 15. The three types of translocation events observed when translocating 10 kbp dsDNA through the nanopore without low-pass filter at 500 kHz. Type 1 is the predominant one where it consists of a single drop, type 2 is where it contains a blip, type 3 is when the event contains two current increase level.

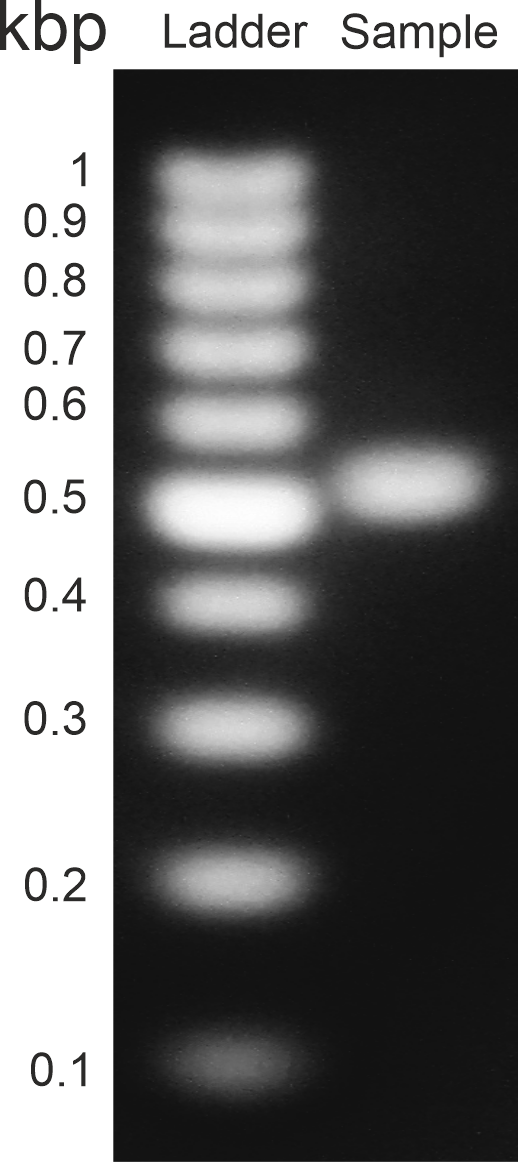

Supporting Figure 16. Gel electrophoresis analysis of the gel extracted 500 bp dsDNA. 1 µg of the gel extracted 500 bp dsDNA was analysed by gel electrophoresis. A DNA ladder was run alongside. The extracted sample migrated at around 500 bp.

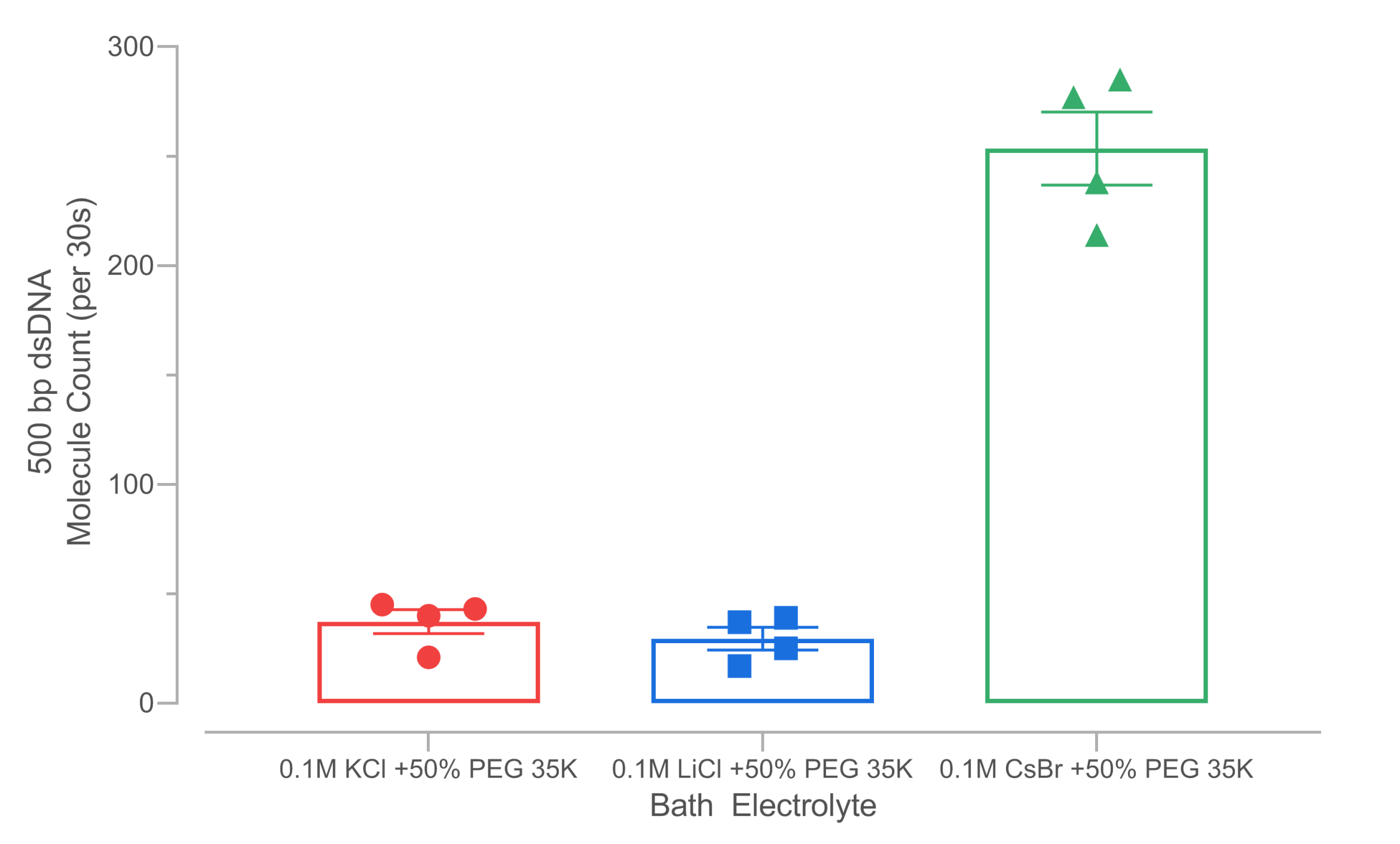

Supporting Figure 17. The molecule count of the translocation of 500 bp dsDNA in different PEG crowded salt bath. The CsBr has the highest current magnitude signal-to-noise ratio and consequently the number of observed events were the highest. N = 4. Error bars are S.E.M.

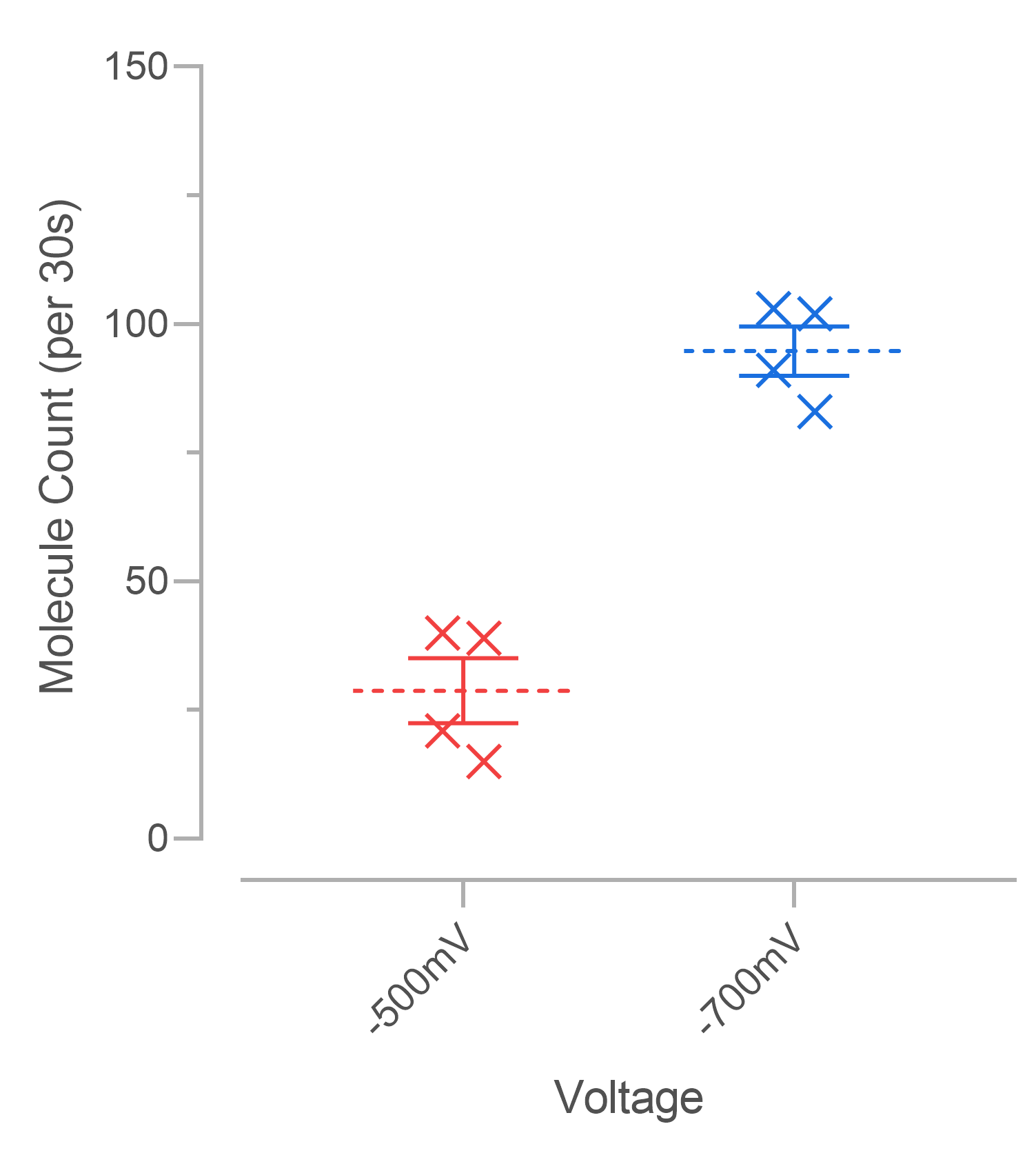

Supporting Figure 18. The translocation molecule count under different low pass filter bandwidth. Error bars are S.E.M.

### Methods

#### Electrolyte bath preparation

Below shows a list of chemicals related to the generation of the electrolyte bath.

| Chemical | Catalogue | Supplier |
| --- | --- | --- |
| Sodium Fluoride (NaF) | 11439933 | Fisher Scientific |
| Potassium Fluoride (KF) | 10318390 | Fisher Scientific |
| Caesium Fluoride (CsF) | 012885 | Alfa Aesar |
| Lithium Chloride (LiCl) | CHE2360 | Scientific Laboratory Supplies |
| Sodium Chloride (NaCl) | S/3160/53 | Fisher Scientific |
| Potassium Chloride (KCl) | P/4240/60 | Fisher Scientific |
| Caesium Chloride (CsCl) | C3032 | Sigma Aldrich |
| Lithium Bromide (LiBr) | 013408 | Alfa Aesar |
| Sodium Bromide (NaBr) | 793574 | Sigma Aldrich |
| Potassium Bromide (KBr) | A16339 | Alfa Aesar |
| Caesium Bromide (CsBr) | 012928 | Alfa Aesar |
| Lithium Iodide (LiI) | 044159 | Alfa Aesar |
| Sodium Iodide (NaI) | 10032620 | Fisher Scientific |
| Potassium Iodide (KI) | PHR1360 | Sigma Aldrich |
| Caesium Iodide (CsI) | 11954211 | Fisher Scientific |
| Dulbecco’s Phosphate buffered saline (PBS) | D8537 | Sigma Aldrich |
| Poly(ethylene) Glycol 4,000 (PEG 4K) | A16151 | Alfa Aesar |
| Poly(ethylene) Glycol 8,000 (PEG 8K) | 043443 | Alfa Aesar |
| Poly(ethylene) Glycol 12,000 (PEG 12K) | 042635 | Alfa Aesar |
| Poly(ethylene) Glycol 35,000 (PEG 35K) | 94646 | Sigma Aldrich |

To generate the electrolyte bath, a similar process as previous publication was used [4]. The alkali metal halide salt was dissolved to 1M with 18.2MΩ ddH_2_O, the solution was then filtered through a 0.22 µm syringe membrane filter (E4780-1223; Starlab UK). The 1M salt solution was then diluted to 0.1M salt solution in 18.2MΩ ddH_2_O followed by 0.22 µm filtration to generate the electrolyte bath. To generate 10 ml of the 50% (w/v) poly(ethylene) glycol (PEG), 1 ml of the 0.22 µm filtered 1M salt solution, 4 ml of 0.22 µm filtered 18.2 MΩ ddH_2_O and 5g of PEG were mixed inside a tube. The tube was then left inside a 70°C incubator for 2 hours followed by incubating at 37°C overnight. The tubes were then left on bench for 4 hours to reach the room temperature prior to use. All electrolytes were stored at room temperature.

#### dsDNA preparation

To prepare the 4.8 kbp dsDNA used in this study, the pmCherry-NLS was linearized by restriction digestion using the enzyme BamHI-HF (R3136; New England Biolabs) with the CutSmart Buffer (B7204S; New England Biolabs) according to the manufacturer’s instruction. The pmCherry-NLS was a gift from Martin Offterdinger (Addgene plasmid # 39319; <http://n2t.net/addgene:39319> ; RRID: Addgene_39319) [5]. The linearized DNA was cleaned and concentrated by the Genomic DNA clean and Concentrator kit according to the manufacturer’s instruction (D4010; Zymo Research). All dsDNA was eluted in the Monarch® DNA Elution Buffer (T1016L; new England BioLabs Inc.). The concentrated dsDNA concentration was measured by UV absorbance at 260 nm with a NanoDrop^TM^ 2000 (Thermo Fisher). 1 µg of the linearized plasmids was analysed by gel electrophoresis with a 1% agarose gel with 1X SYBR safe dye (S33102; Thermo Fisher) incorporated into the gel. 1 µg of TrackIt^TM^ 1 Kb Plus DNA Ladder (10488085; Thermo Fisher) was run alongside the dsDNA samples. The dsDNA was kept at -20°C for storage.

To prepare the 10 kbp dsDNA, the GeneRuler 1 kb Plus DNA Ladder (SM1331; Thermo Fisher) was separated via a 0.8% agarose gel, the 10 kbp band were physically isolated with a blade and the 10 kbp dsDNA was extracted using the Monarch® DNA Gel Extraction Kit according to the manu(T1020; New England BioLabs Inc.), the extracted dsDNA was further purified using the Genomic clean and Concentrator kit (D4010; Zymo Research). All dsDNA was eluted in the Monarch® DNA Elution Buffer (T1016L; new England BioLabs Inc.) and stored at -20°C.

The lyophilised 75 bp dsDNA (SM1431; Thermo Fisher) was redissolved in the Monarch® DNA Elution Buffer (T1016L; new England BioLabs Inc.) and stored at -20°C.

All the dsDNA was then diluted from stock to 0.3 nM with 0.1M KCl, 0.1M LiCl or 1XPBS prior to use.

#### Scanning electron microscopy

The nanopores of the nanopipettes were imaged by scanning electron microscopy (Leo 1530 FEG-SEM; Zeiss). Nanopipettes were first sputter coated with a gold layer of a few nanometres in thicknesses. The nanopipettes were then mounted onto the sample holder and tilted to an angle of 60 and above for imaging. The nanopipettes were imaged at between 2 and 3 kV at a working distance of 5 mm and below at an aperture size of 30.00 m using a InLens detector.
